# A unique mechanism of snRNP core assembly

**DOI:** 10.1101/2024.07.09.602679

**Authors:** Yingzhi Wang, Xiaoshuang Chen, Xi Kong, Yunfeng Chen, Zixi Xiang, Yue Xiang, Yan Hu, Yan Hou, Shijie Zhou, Congcong Shen, Li Mu, Dan Su, Rundong Zhang

## Abstract

The assembly of spliceosomal snRNP cores involves seven Sm proteins (D1/D2/F/E/G/D3/B) forming a ring around each snRNA and typically requires several essential assembly chaperones, particularly the SMN-Gemins complex implicated in spinal muscular atrophy (SMA). Strikingly, in budding yeast, snRNP core assembly only involves Brr1, a nonessential homolog of Gemin2. To unravel this enigma, we systematically investigated snRNP core assembly in budding yeast using biochemical and genetic approaches. We discovered two distinct pathways in budding yeast: chaperone-mediated and direct. The chaperone-mediated pathway involves two proteins, Brr1 and a novel protein, Lot5, but is inefficient. Lot5 binds D1/D2/F/E/G to form a heterohexameric ring (6S). Brr1 binds D1/D2/F/E/G and 6S but cannot displace Lot5 to facilitate assembly. Disruption of both BRR1 and LOT5 genes caused mild growth retardation, but LOT5 overexpression substantially impeded growth. The direct pathway uniquely involves F/E/G as a trimer and a stable D1/D2/F/E/G intermediate complex, explaining the non-essentiality of assembly chaperones. These findings unveil a unique assembly mechanism of snRNP cores, illuminate the evolution of assembly chaperones, and suggest avenues for studying SMA pathophysiology.

## INTRODUCTION

U-rich small nuclear ribonucleoprotein particles (snRNPs) are essential building blocks for the spliceosomes, which are responsible for removing introns and splicing exons in eukaryotic pre-mRNAs ^1, 2^. Excluding U6 and U6atac snRNPs, U-rich snRNPs (including U1, U2, U4 and U5 for the major spliceosome, and U11, U12, U4atac and U5 for the minor spliceosome) have a core structure in common. This core structure is composed of a single U-rich snRNA and seven different Sm proteins, forming a ring-like structure, arranged in the order of D1-D2-F-E-G-D3-B around a specific sequence in the snRNA termed the Sm site, which is typically PuAUUUNUGPu (Pu=purine) ^1, 2^. Sm proteins belong to a protein family that includes Sm-like (Lsm) proteins, and they share a common Sm fold characterized by an N-terminal helix followed by 5 highly bent β-strands ^3^. U6 and U6atac snRNPs also have similar core structures around the 3’-ends of their snRNAs, but they contain a different set of Sm-like proteins known as Lsm2-8 ^1, 2^.

The biogenesis of the Sm class snRNPs is a complex process. The snRNAs are transcribed in the nucleus and then transported to the cytoplasm. In the cytoplasm, the seven Sm proteins assemble on the snRNAs, and their 5’-caps undergo hypermethylation. Subsequently, the snRNPs are imported back into the nucleus where further modifications occur, and additional proteins are assembled. The biogenesis process occurs in several phases and involves numerous interacting proteins and assembly chaperones ^1,4^. In contrast, Lsm class snRNPs are entirely assembled within the nucleus and do not require the assistance of chaperones for their formation ^1^.

The formation of snRNP cores—the assembly of seven Sm proteins onto the Sm site of snRNA— occurs in the cytoplasm and requires the assistance of several assembly chaperones ^1, 4, 5^. Interestingly, early studies have demonstrated that the snRNP cores can spontaneously assemble by mixing snRNA and three existing Sm subcomplexes, namely SmD1/D2, SmF/E/G, and SmD3/B ^6^. It has also been established that the assembly order of the Sm core proceeds through a stable SmD1/D2/F/E/G–snRNA intermediate, referred to as the Sm subcore, and eventually maturates by recruiting SmD3/B ^6^. In most eukaryotes, the assembly of the Sm core requires assistant proteins that are organized into two sets of protein complexes—the PRMT5 (protein arginine methyltransferase 5) complex and the SMN (survival of motor neuron) complex ^1, 4, 5^. In higher eukaryotes such as vertebrates, the PRMT5 complex consists of three proteins: PRMT5, WD45 and pICln. On the other hand, the SMN complex is composed of nine proteins: SMN, Gemin2-Gemin8, and Unrip. Even in less complex unicellular eukaryotes like *Schizosaccharomyces pombe*, there are also many orthologues of these two complexes, including pICln, SMN, Gemin2 and Gemin6-8 ^7, 8, 9, 10, 11, 12^. Many of these proteins are essential, and their individual deletion, such as SMN, Gemin2, and pICln, results in early embryonic death in vertebrates ^13, 14, 15^. In *S. pombe*, the deletion of any of the members of the SMN complex, including SMN, Gemin2 and Gemin6-8, hampers cell growth ^7, 8, 9, 10, 12^. Deficiencies in SMN expression cause spinal muscular atrophy (SMA), a devastating motor neuron degenerative disease, with an incidence frequency of about 1/10,000 in humans ^16, 17^. Some orthologous proteins in *S. pombe*, such as pICln, despite being nonessential, play an important role in optimal yeast cell growth ^11^. Therefore, many of these chaperone proteins have essential functions in the assembly of snRNP cores.

Extensive biochemical and structural studies have provided insights into the mechanism of Sm core assembly facilitated by the two chaperone complexes. These complexes play distinct roles in two consecutive phases of the assembly process. In the first phase, pICln, a protein predominantly comprised of a pleckstrin homology (PH) domain fold that shares similarity with the Sm fold, recruits SmD1/D2 and SmF/E/G to form a ring-shaped 6S complex. This complex stabilizes the five Sm proteins (5Sm) in the appropriate assembled order and simultaneously prevents RNA binding to 5Sm ^5, 18^. Additionally, pICln also binds SmD3/B ^5^. The PRMT5/WD45 complex methylates the C-terminal arginine residues of SmB, SmD1 and SmD3, which is believed to enhance the interactions between the Sm proteins and SMN. However, in certain species, the C-terminal methylation of these Sm proteins is not necessary ^19^.

In the second phase, the SMN complex accepts 5Sm and releases pICln ^5, 12^. Gemin2, a component of the SMN complex, is the acceptor of 5Sm ^20^. Gemin2 has two structural domains: an N-terminal domain (ND) containing an α-helix followed by a short β-strand, and a C-terminal domain (CD) consisting of a helical bundle. These domains are connected by a flexible loop ^20^. Gemin2 binds to the outer perimeter of the horseshoe-shaped 5Sm using its ND to interact with SmF/E and its CD to interact with SmD1/D2. In addition, the CD of Gemin2 also binds to the N-terminal helix of SMN (residues 26-51 in humans) via a different surface ^20^. Apart from interacting with Gemin2 through its N-terminal domain, SMN also interacts with Gemin8 through its C-terminal self-oligomerized YG box ^21^. Gemin8 further binds Gemin6/7 and Unrip, although the specific roles of these interactions are not well understood ^21, 22, 23^. Gemin3 contains a DEAD box domain and is a putative RNA helicase ^24^. Gemin4 usually forms a complex with Gemin3, but its role remains unknown ^25^. Gemin5 was identified to initially bind pre-snRNAs and deliver them to the rest of the SMN complex for assembly into the Sm core ^26^. It also plays additional roles such as cooperating in translation control and ribosome binding ^27^. However, Gemin3-5 and Unrip are not found in unicellular eukaryotes such as *S. pombe*, implying that they may not be as essential as the other members ^10, 12^.

Despite the conservation of many components in both complexes, there are exceptions to their essentiality. For example, the orthologues of Gemin2 and pICln in *Arabidopsis thaliana* have been found to be nonessential ^28, 29^. Another instance is observed in the budding yeast *Saccharomyces cerevisiae*, where Brr1, a weak homolog of Gemin2, is found to play a role in Sm core assembly; however, it is also nonessential ^30, 31^. Although *S. cerevisiae* contains a lower percentage of introns in its genome than most other eukaryotes ^32^, disruption of any of the seven Sm genes is not tolerated ^33^. This indicates the indispensability of these Sm proteins and the Sm core assembly process. Furthermore, similar to vertebrate snRNAs, yeast pre-snRNAs are reported to be exported to the cytoplasm for Sm core assembly, which is vital for the re-import of snRNAs into the nucleus ^34^. Hence, numerous puzzles still remain unanswered. Why do the essential assembly chaperones in most other eukaryotes become nonessential in *Arabidopsis* and *S. cerevisiae*? Do they have distinct chaperones and unique mechanisms in their snRNP core assembly? If so, does this divergence provide any insights into the evolutionary aspects of the assembly chaperone system in snRNP core assembly?

To address these questions, the current study focused on investigating the assembly chaperone system involved in Sm core formation in *S. cerevisiae*. We identified Lot5, a protein with an unknown function, as being involved in Sm core assembly. We characterized Lot5, along with the weak Gemin2 homologue Brr1, both biochemically and genetically in the context of Sm core assembly. This study uncovered a unique mechanism of Sm core assembly in *S. cerevisiae*, shedding light on the evolution of assembly chaperones in eukaryotes. These findings have potential implications for studying SMA biology and pathophysiology.

## RESULTS

### Identification of Lot5 as a protein associated with Sm proteins and Brr1 in *S. cerevisiae*

Thus far, only Brr1, the putative orthologue of Gemin2, has been identified and partially characterized in *S. cerevisiae* to play a role in the production of spliceosomal snRNPs ^30, 31^. Given that most other eukaryotes have a more elaborate assembly chaperone system for the Sm core, incorporating as many as six proteins (SMN, Gemin2, Gemin6-8 and pICln) even in the unicellular *S. pombe* ^10, 12^, it is speculated that there could be additional components beyond Brr1 involved in Sm core assembly in *S. cerevisiae*. Using a bioinformatics approach, we identified Lot5 as a potential orthologue of pICln (12.7% identity and 24.5% similarity to human pICln, and 13.1% identity and 28.8% similarity to *S. pombe* pICln). Although Lot5 shares a low sequence similarity with pIClns in other species, the secondary structure prediction indicates that like pIClns, it also has a PH domain fold. More significantly, regions of Lot5 predicted to interact with Sm proteins—specifically the β5 strand and α-helix—display a high degree of similarity to those of pICln from other species (Fig. 1A). Additionally, the AlphaFold structural model of Lot5 (Fig. S1) aligns well with the crystal structure of pICln, although it contains extended loops at several junctures between secondary structure elements, including loops L0, L1, L5, and L6 (Fig. 1B). To confirm whether Lot5 is a pICln orthologue and to identify potential proteins associated with Sm core assembly, we constructed plasmids containing Lot5, Brr1 or EGFP (as a negative control) with an N-terminal tandem affinity purification (TAP) tag and transformed each of them into a budding yeast strain. After conducting two steps of affinity purification on the extracts from the yeast transformants, the pulled-down samples were subjected to SDS-PAGE followed by Western blot analysis using monoclonal anti-FLAG antibody as well as silver staining (Fig. 1C-D). The TAP-tagged EGFP and Brr1 were clearly identified as single bands, whereas the TAP-tagged Lot5 appeared in multiple bands, indicating C-terminal degradations (Fig. 1C). The purified protein samples were further subjected to mass spectrometry analysis (Table S1). The Brr1 pull-down sample showed significant identification of Sm proteins, as anticipated. However, Lot5 was also significantly identified in the Brr1 pull-down sample. In the proteins enriched by Lot5, Brr1 was significantly identified, and several Sm proteins were also identified in the Lot5 pull-down sample. These results suggest that Lot5 is associated with Sm proteins and Brr1, and is likely involved in Sm core assembly.

**Fig. 1.**
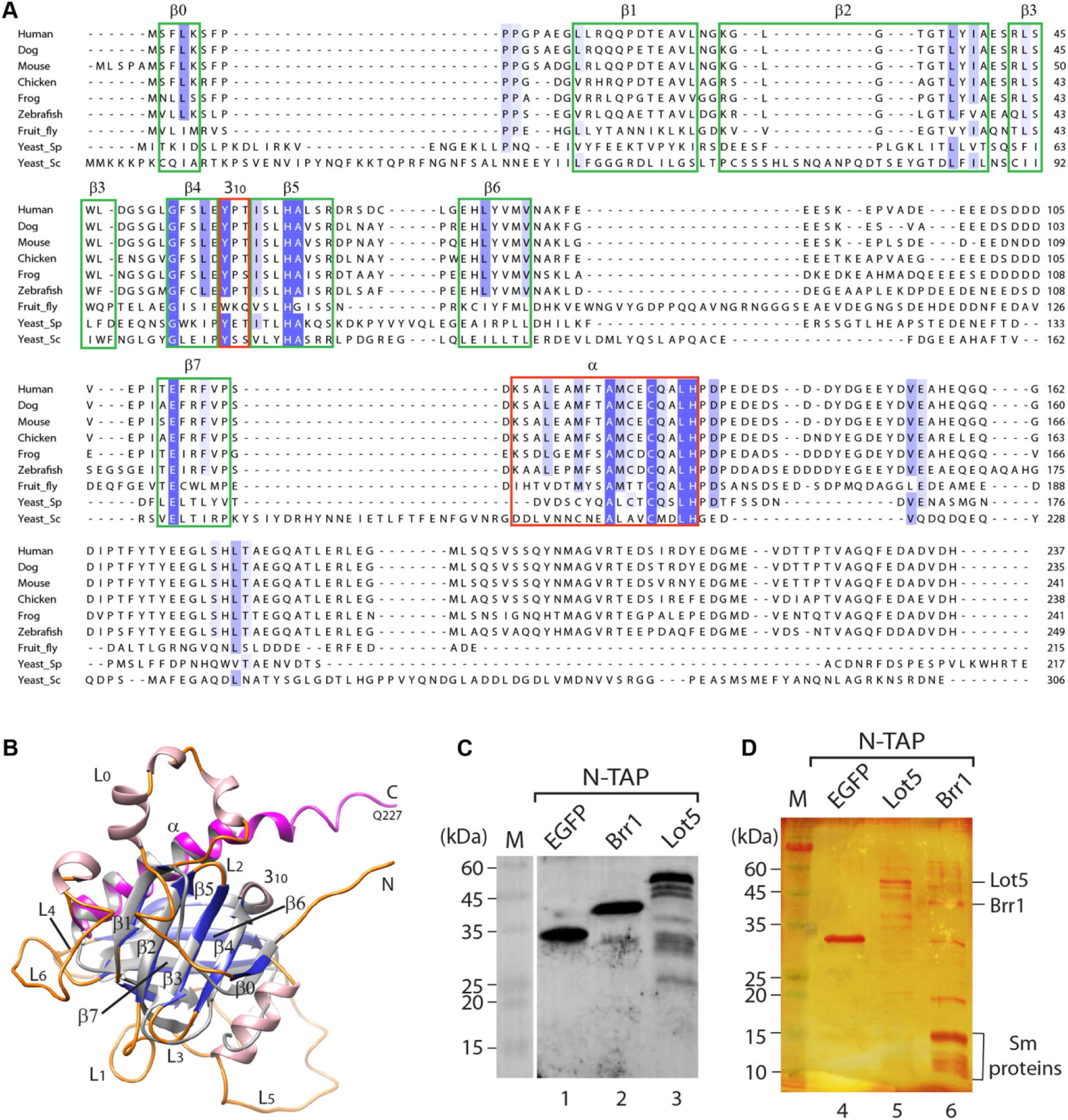
Alignment of Lot5 with pICln homologs and identification of proteins interacting with Lot5 and Brr1. (A) Sequence alignment of Lot5 (Yeast_Sc) with pICln homologs from various species. Secondary structures (β0-β7 strands, 3_10_ and α helices) are annotated based on the crystal structure of fruit fly pICln (PDB: 4F7U). Conserved residues are highlighted in blue, with the intensity of the color reflecting the level of conservation. (B) Structural alignment between the AlphaFold structure model of Lot5 (residues 1-227) (β0-β7 strands are colored in blue, α helix in purple, other α helices in pink, and loops in orange) and the structure of fruit fly pICln (in grey); (C-D) Identification of proteins interacting with the N-terminally TAP-tagged Lot5 or Brr1 by a pull-down assay followed by Western blot analysis probed with a FLAG-antibody (C) and silver staining (D), as well as mass spectrometry analysis (Table S1). EGFP served as a control.

### Distinct Sm subcomplex interactions

To further characterize the interactions among Lot5, Brr1 and Sm proteins *in vitro*, we constructed expression plasmids containing these proteins and expressed them in *E. coli*. Initially, we aimed at characterizing the interactions among the Sm proteins from budding yeast *in vitro*, which has never been undertaken before. Similar to making human Sm proteins expression ^20, 35^, we co-expressed SmD1/D2, SmF/E/G and SmD3/B in their putative subcomplex forms respectively. Indeed, the Sm proteins in the three Sm subcomplexes were successfully purified together as evidenced by the purification process (Fig. 2). We next examined the interactions between Sm subcomplexes using Ni-beads pull-down assay. Immobilized D1/D2 pulled down F/E/G, but not D3/B, even when all three Sm subcomplexes were present within the mixture (Fig. 2A). Consistently, Immobilized F/E/G pulled down D1/D2, but not D3/B (Fig. 2B).

**Fig. 2.**
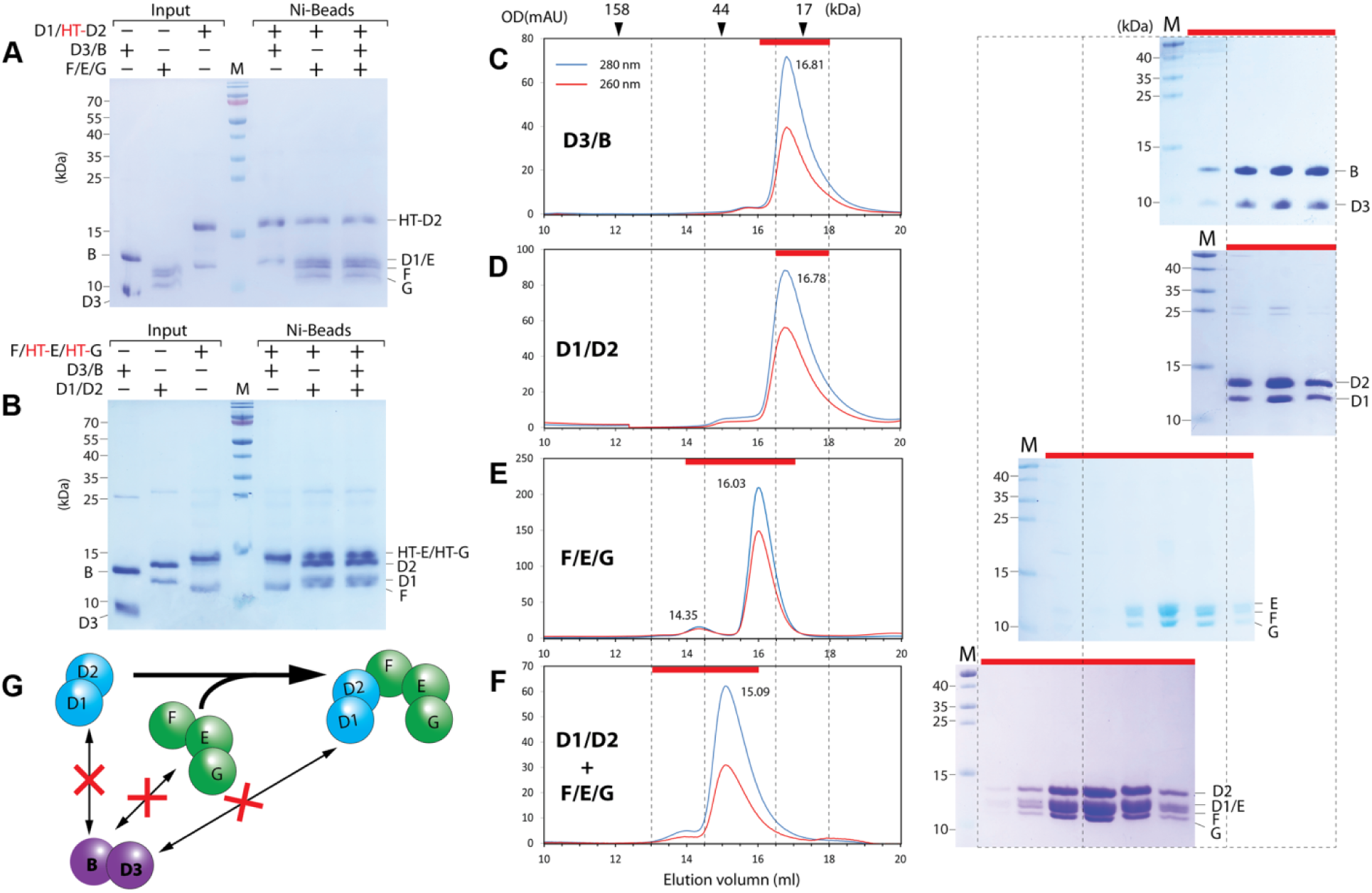
Oligomerization states of 3 Sm subcomplexes from *S. cerevisiae* and their interactions. (A-B) Interactions among Sm subcomplexes were tested by a pull-down assay. In each reaction, only one Sm subcomplex, D1/D2 (A) or F/E/G (B), was tagged with His-TEV (HT-), while the other subcomplex(es) had the HT-tag removed. The indicated Sm subcomplexes were incubated and then pulled down using Ni-beads. M: markers. (C-F) The oligomerization states of D3/B (C), D1/D2 (D), and F/E/G (E) and the interaction between D1/D2 and F/E/G (F) were tested using GFC (left panels). The eluted fractions (indicated by red bars) were then analyzed by SDS-PAGE/CBB staining (right panels). For each set, a representative result from two independent experiments is shown. (G) Summary of the oligomerization states and interactions of the three Sm subcomplexes.

We further used gel filtration chromatography (GFC) to study the oligomerization state of each of the three Sm subcomplexes and their complexations. As expected, both D3/B [Fig. 2C, peak at 16.81 ml, calculated apparent molecular weight (aMW) =17.8 kDa] and D1/D2 (Fig. 2D, peak at 16.78 ml, aMW =18.3 kDa) were eluted as single dimers. Surprisingly, F/E/G was predominantly eluted as a trimer (Fig. 2E, peak at 16.03 ml, aMW =27.2 kDa, ∼93% of the total F/E/G estimated using the area under curve). Although there was a tiny elution peak at 14.35 ml (∼7%), the peak faction was hardly visible on SDS-PAGE followed by Coomassie bright blue (CBB) staining. This finding contrasts significantly with observations from human ^5, 35^ and *S. pombe* (*Sp*) F/E/G ^12^. Human F/E/G have been known to form a hexamer ^6^, although it still has a small fraction of trimer (10-40%) when characterized by GFC ^5, 35^, while *Sp-*F/E/G was completely a hexamer by GFC ^12^.

The mixture of D1/D2 and F/E/G was eluted with a single peak at 15.09 ml (Fig. 2F, aMW= 44.8 kDa), which is distinguished from the two input subcomplexes alone (Fig. 2, D-E), and the SDS-PAGE/CBB staining showed that the peak contained the bands of D1/D2 and F/E/G with equal stoichiometric ratios. This indicates that D1/D2 and F/E/G can form a stable D1/D2/F/E/G pentamer (5Sm). This observation is also significantly distinct from the human and *S. pombe* versions of D1/D2 and F/E/G. Although a mixture of human D1/D2 and F/E/G could form a substantial portion of D1/D2/F/E/G pentamers, a significant portion of D1/D2 and F/E/G still existed alone in a GFC assay ^35^. The mixture of *Sp*-D1/D2 and *Sp*-F/E/G cannot form D1/D2/F/E/G pentamer at all ^12^. When all three Sm subcomplexes were incubated and analyzed by GFC, they were eluted with two peaks at positions similar to the 5Sm complex and D3/B alone (Fig. S2), indicating that there is little cooperativity among them to form a larger oligomeric form. The results are summarized in Fig. 2G.

We further tested the binding of 5Sm with a customized minimal snRNA, Umini-snRNA, which contained the Sm site followed by a stem-loop [this feature has been termed as snRNP code, fully accounting for Sm core formation ^35, 36^], or 5Sm and D3/B with Umini-snRNA, using either electrophoresis mobility shift assay (EMSA) or GFC approaches, and found that both could specifically bind to Umini-snRNA, but not to Umini-ΔSm-snRNA, in which “UUUUU” of the Sm site “AGUUUUUGA” was replaced by “CCCCC” (Fig. 3G). Stable Sm subcore or Sm core could form (Fig. S2). In addition, as this customized snRNA behaves similarly to U4-snRNA in Sm core assembly (compare Fig. 3G with Fig. S3), it would be used in the characterization of the assembly process involving Lot5 and Brr1.

**Fig. 3.**
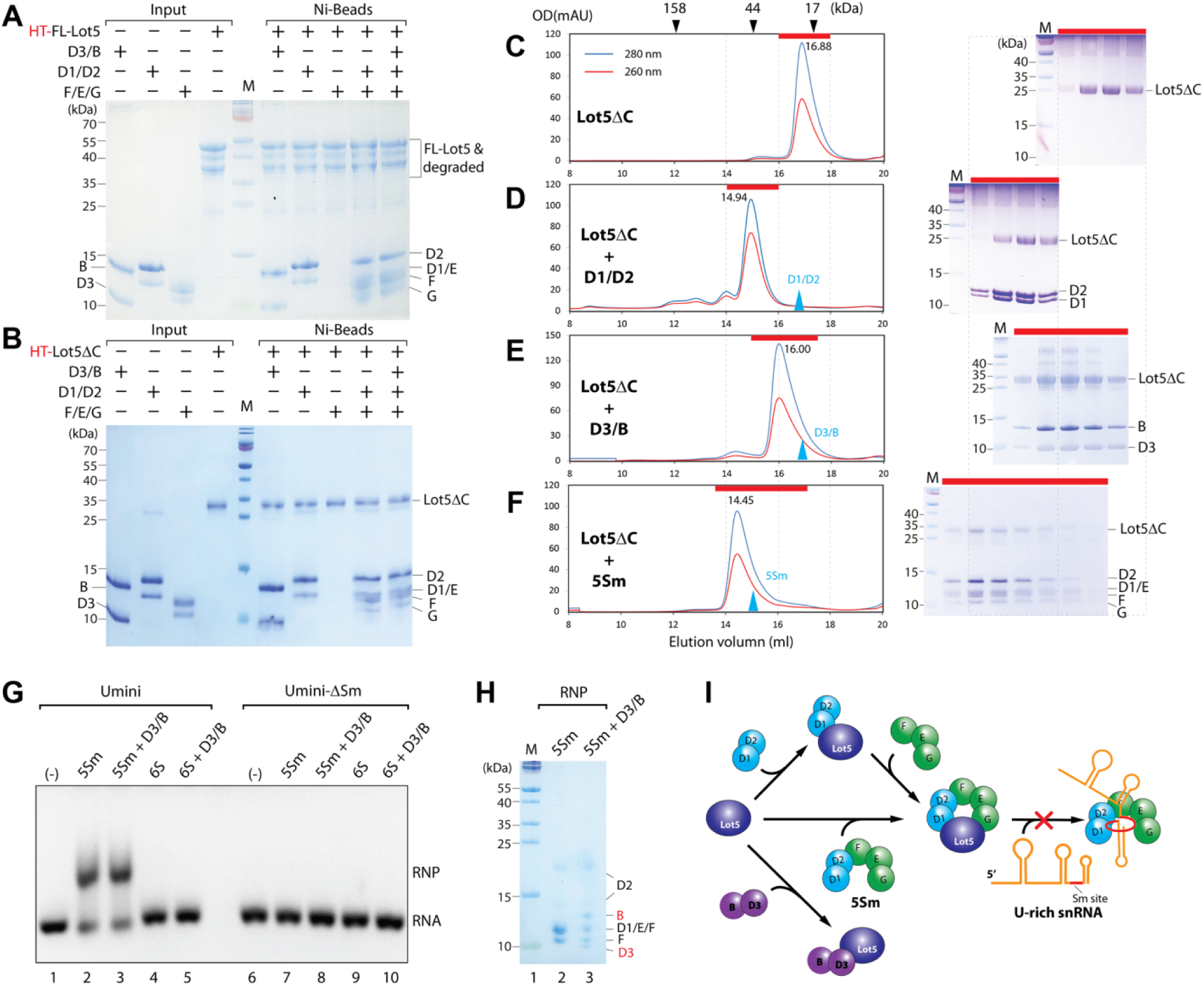
Lot5 acts as a pICln homolog in interacting with the Sm subcomplexes and snRNA. (A-B) The binding assay was performed using full-length Lot5 from *S. cerevisiae* (A) or a C-terminally truncated form of Lot5 (B), with each of the three Sm subcomplexes individually and together. The interaction was assessed through a pull-down assay using Ni-beads. M: markers. (C-F) Complexation assays. Lot5 alone (C) or equimolar amounts of Lot5 mixed with D1/D2 (D), D3/B (E), or 5Sm (F), was subjected to GFC (left panels). The eluted fractions (indicated by red bars) were then analyzed by SDS-PAGE/CBB staining (right panels). The blue arrowheads indicate the GFC peaks of the input components (D1/D2, D3/B and 5Sm) for comparison. (G) Lot5 blocks snRNA binding to the Sm subcore and core. Reconstituted 5Sm or 6S, either alone or together with D3/B, was pre-incubated with Umini-snRNA or Umini-ΔSm-snRNA (as a control) and subjected to electrophoresis mobility shift assay (EMSA). (H) The protein components binding to Umini-snRNA, of the shifted bands in panel (G), were analyzed using SDS-PAGE/CBB staining. For each set, one representative of at least two independent experiments is shown. (I) A cartoon model of the interactions of Lot5 with the Sm proteins and snRNA.

### Lot5 acts as pICln in interacting with Sm subcomplexes and U-snRNA

As Lot5 contains a longer C-terminus that is likely sensitive to degradation, and truncations of the C-terminal sequences did not affect the function of pICln in other species, we expressed both the full-length Lot5 (FL-Lot5) and a C-terminal truncated version of Lot5, Lot5 (residues 1-227) or Lot5ΔC, from *E. coli* for pull-down interaction study with Sm proteins. As anticipated, FL-Lot5 did degrade significantly (Fig. 3A). Nevertheless, both FL-Lot5 and Lot5ΔC interacted with the 3 Sm subcomplexes in the same pattern (Fig. 3, A-B). When mixed with a single Sm subcomplex, both pulled down D3/B or D1/D2, but not F/E/G. When mixed with D1/D2 and F/E/G, both pulled them down together. When mixed with D1/D2 and F/E/G for a period of time, and then mixed with D3/B, both only pulled down D1/D2 and F/E/G, with D3/B hardly detected. Because Lot5ΔC interacts with Sm proteins in the same way as FL-Lot5 and does not degrade (Fig. 3B), we used it for further *in vitro* characterization.

We further examined the complexation of Lot5ΔC and Lot5ΔC-Sm subcomplexes using GFC. Lot5ΔC alone was eluted in a single peak at 16.88 ml, corresponding to aMW of 17.1 kDa (Fig. 3C), indicating that it is a monomer. The mixture of Lot5ΔC and D1/D2 had a peak eluted at 14.94 ml (aMW=48.6 kDa), which is distinct from the peaks of Lot5ΔC and D1/D2 alone (Fig. 3D), indicating the formation of the Lot5ΔC/D1/D2 heterotrimer. Similarly, Lot5ΔC and D3/B also had an eluted peak (16.00 ml, aMW = 27.5 kDa) distinct from Lot5ΔC and D3/B alone (Fig. 3E), indicating a heterotrimeric state of Lot5ΔC/D3/B. When Lot5ΔC was mixed with 5Sm, the elution peak (14.45 ml, aMW= 63.2 kDa) appeared earlier than those of Lot5ΔC and 5Sm alone (Fig. 3F), indicating the formation of a heterohexamer of Lot5ΔC/5Sm (referred to as 6S). When 6S was mixed with D3/B, the elution peaks remained the same as those of 6S and D3/B alone (Fig. S4D), indicating that D3/B cannot displace Lot5ΔC from 6S, which is consistent with the pull-down observations (Fig. 3B). These interactions with Sm proteins exhibit similar behavior to those of human pICln and *Sp*-ICln that were previously characterized ^5, 12^.

In humans, pICln binds D1/D2 and F/E/G to form a heterohexameric ring, which prevents snRNA from binding to 5Sm and forming a subcore ^5, 18^. To test if this is also the case for Lot5ΔC, we performed EMSA using Umini-snRNA. As positive controls, 5Sm or the mixture of 5Sm and D3/B bound Umini-snRNA and caused a shift in the RNA band (Fig. 3G&H, lanes 2-3). The specificity of Sm subcore or core assembly was demonstrated by the fact that the negative RNA control, Umini-ΔSm-snRNA, could not bind any of the protein complexes mentioned above (Fig. 3G, lanes 7-8). In contrast, 6S alone or in mixture with D3/B did not cause a significant shift in the RNA band (Fig. 3G, lanes 4-5). The inability of 6S alone or in mixture with D3/B to bind Umini-snRNA was further confirmed by GFC (Fig. S4). Furthermore, the inability of 6S alone or in mixture with D3/B to bind U4-snRNA was confirmed by EMSA (Fig. S3). These results further indicate that Lot5 behaves similarly to pICln and the interactions of Lot5 with Sm proteins and U-rich snRNA are summarized in Fig. 3I.

### Brr1 functions as Gemin2/SMN_Ge2BD_ in Sm core assembly

Although Brr1 has been biochemically characterized to play a role in the manufacture of spliceosomal snRNPs ^31^ and to interact with Sm proteins genetically ^30^, a direct interaction between reconstituted Brr1 and Sm proteins has not been demonstrated. Therefore, we performed a Ni-beads pull-down assay to examine the interactions between Brr1and Sm subcomplexes (Fig. 4A). Brr1 was found to pull down F/E/G significantly but could not pull down D3/B or D1/D2. In addition, Brr1 pulled down D1/D2 and F/E/G when incubated with both, but it only pulled down D1/D2 and F/E/G when all the 3 Sm subcomplexes were tested together. We also analyzed the oligomeric state of the complexes using GFC. Brr1 was eluted at 15.09 ml (Fig. 4B), which corresponds to aMW of 44.8 kDa, indicating that Brr1 exists as a monomer. The mixture of Brr1 and F/E/G was eluted at 3 peaks (Fig. 4C), which were identified as Brr1/F/E/G, Brr1 and F/E/G respectively. The presence of both the input components and the final complex in the elution peaks suggests that the interaction between Brr1 and F/E/G is in equilibrium. When the mixture of Brr1 and 5Sm was analyzed by GFC, only a single peak was eluted at 13.54 ml (aMW=102.9 kDa), which contained all the proteins with stoichiometric ratios, indicating that Brr1 and 5Sm form a stable heterohexamer (Fig. 4D). This differs significantly from the Gemin2 orthologues in other species. For example, *Sp*Gemin2 from *S. pombe* cannot form a stable complex with D1/D2 and F/E/G without the Gemin2-binding domain (Ge2BD) of *Sp*SMN (residues 1-35) bound, but *Sp*SMN(1-35)/SpGemin2 can ^12^. This indicates that the single protein Brr1 acts as a combination of Gemin2 and the Ge2BD of SMN. Moreover, Brr1 does not contain any YG-box oligomerization domain at the C-terminus of SMN because Brr1 exists as a monomer.

**Fig. 4.**
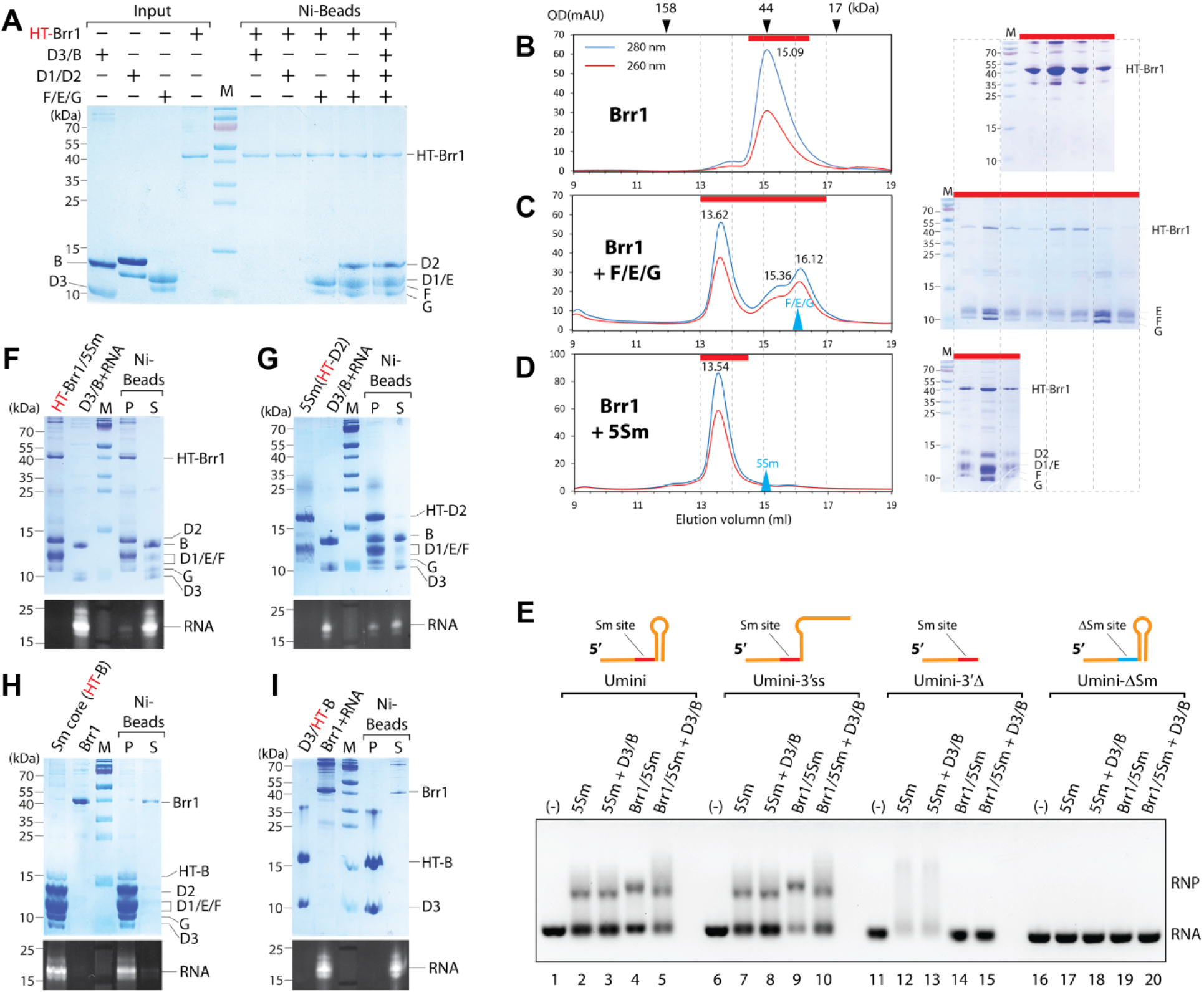
Brr1 acts like Gemin2/SMN_Ge2BD_ in interacting with the Sm subcomplexes and snRNA. (A) The binding assay was performed using *S. cerevisiae* Brr1 with each of the three Sm subcomplexes individually and together. The interaction was assessed through a pull-down assay using Ni-beads. (B-D) Complexation assays. Brr1 alone (B), or equimolar amounts of Brr1 mixed with F/E/G (C) or 5Sm (D), were subjected to GFC (left panels). The eluted fractions (red bar) were analyzed by SDS-PAGE/CBB staining (right panels). Blue arrowheads indicate the GFC peaks of the input components (F/E/G and 5Sm) for comparison. (E) The role of Brr1 in assembly of the Sm subcore and core was assessed using EMSA. Reconstituted 5Sm or Brr1/5Sm, either alone or together with D3/B, was pre-incubated with Umini, Umini-3’ss, Umini-3’Δ, or Umini-ΔSm snRNA and subjected to EMSA. One representative of two independent experiments is shown. (F-I) Ni-beads pull-down assays were performed to test the dissociation of Brr1 from Sm core assembly. The forward reaction was initiated with either HT-Brr1/5Sm (F) or 5Sm (HT-D2) as a positive control (G) and incubated with D3/B and Umini-snRNA. The reverse reaction started with pre-assembled Sm core (HT-B) (H) or HT-B/D3 plus Umini-snRNA (I) and incubated with Brr1. The samples were then analyzed using SDS-PAGE /CBB staining and GelRed staining.

As previously reported in our studies, both human and *S. pombe* Gemin2/SMN_Ge2BD_ can bind to D1/D2/F/E/G to form a stable intermediate complex known as 7S. The 7S complex can accept the cognate snRNAs and D3/B to form Sm cores and simultaneously release Gemin2/SMN_Ge2BD_ ^12, 35^. We expect the same behavior for the Brr1/5Sm complex. In EMSA, Brr1/5Sm or Brr1/5Sm plus D3/B caused a shift in the Umini-snRNA band, similar to 5Sm or 5Sm plus D3/B (Fig. 4E, lanes 1-5). In contrast, they did not cause a shift in any of Umini-ΔSm-snRNA bands (Fig. 4E, lanes 16-20). This indicated that Brr1/5Sm specifically binds to snRNAs. Moreover, Brr1/5Sm or Brr1/5Sm plus D3/B also caused a shift in the bands of U4-snRNA (Fig. S3). In addition, the selectivity of 5Sm bound by Brr1 was tested on two derivatives of Umini-snRNA: Umini-3’ss, which changed 3’SL into a linear single strand, and Umini-3’Δ, which deleted 3’SL (Fig. 4E, lanes 6-15). While Brr1/5Sm or Brr1/5Sm plus D3/B caused a shift the bands of Umini-3’ss, they could not shift Umini-3’Δ. These results indicated that, like Gemin2 in humans ^35^, the binding of Brr1 to 5Sm enhances the selectivity of RNA for Sm core assembly.

To test if the formation of the Sm core leads to the release of Brr1, we used reconstituted HT-Brr1/5Sm to incubate with D3/B and Umini-snRNA to allow the assembly reaction to occur. We then used Ni-beads to separate the fractions bound to HT-Brr1 and those that were unbound (Fig. 4F). In the precipitated fraction, 5Sm were still bound to HT-Brr1, but only a tiny amount of Umini-snRNA was visible and D3/B was completely absent. In the supernatant fraction, most of 5Sm were visible along with D3/B and Umini-snRNA. In the control experiment, 5Sm (with HT-tag on SmD2) without Brr1 bound was able to pull down Umini-snRNA and D3/B significantly, indicating the formation of the Sm core (Fig. 4G). The mutual exclusion between D3/B and Brr1 in the Sm core formation indicates the release of Brr1 from mature Sm core, which is consistent with the negative cooperativity mechanism between Gemin2/ SMN_Ge2BD_ and RNA in binding to 5Sm ^35^. We also used reconstituted Sm core (with HT-tag on SmB) to incubate with Brr1 and then used Ni-beads to pull down the Sm core to test if Brr1 binds to it (Fig. 4H). As expected, Brr1 was not pulled down by the Sm core, which further confirmed that the formation of the Sm core decreases the binding affinity between Brr1 and 5Sm in the reverse direction. In another control experiment, HT-B/D3 could not pull down Umini-snRNA or Brr1 (Fig. 4I). These data further suggest that Brr1 functions as Gemin2/SMN_Ge2BD_ in Sm core assembly process.

### Brr1 contains an extra helical domain which can be removed

Brr1 is longer than human Gemin2 and *Sp*Gemin2, with 341 residues compared to 280 and 235 residues, respectively. The N-terminal E/F-binding domain and the last five α-helices plus a short 3_10_-helix are more conserved within the sequence of Brr1 (Fig. 5A & S5). Secondary structure prediction indicates that Brr1 contains several additional helices, one of which may serve as a substitute for the single helical form of SMN_Ge2BD_. We attempted to purify Brr1-CD (residues 80-341) and reconstitute Brr1/5Sm for crystal structure studies, but were unsuccessful. Recently, more accurate structure prediction programs based on artificial intelligence (AI) such as AlphaFold (AF) and RosettaFold (RF) have been developed ^37,38^. The structural models of Brr1-CD predicted by both programs are similar in the conserved region (residues 201-341), where a very long loop containing a short α-helix (named αE) occupies the position of SMN_Ge2BD_ (Fig. 5, A-B,D). However, the structure models differ significantly in the less conserved region (residues 80-200). Although both models predicted 6-7 helices in this region, the positions of these helices are dramatically different (Fig. 5B and S1). In the AF model, region 80-108 forms a helix that is perpendicular to the last 4 α-helices and interacts with α6 and α8. Region 115-198 contains 5 α-helices, which form a separate helical domain. In the RF model, region 80-150 forms a separate helical domain. Region 166-180 is a helix that occupies the same position as the region 80-108 in the AF model. Region 153-164 and region 184-195 are two small connecting helices. To determine which model is more accurate, we constructed two truncated versions of Brr1-CD: Brr1CDΔ(122-197), which removes the AF-predicted separate domain, and Brr1CDΔ(80-165), which removed the RF-predicted separate domain and the small connecting helix (Fig. 5C). We hypothesized that the construct with higher expression would correspond to the more accurate model. Brr1CDΔ(122-197) was successfully purified at the expected size and detected by anti-His antibody, while Brr1CDΔ(80-165) was poorly purified and none of the sample fractions could be detected by anti-His antibody (Fig. 5C and S6). This indicates that the AF model of Brr1 is more accurate. The well-expression of the C-terminal extra helical domain (EHD) of Brr1 (116-197) also supports this argument (Fig. S6).

**Fig. 5.**
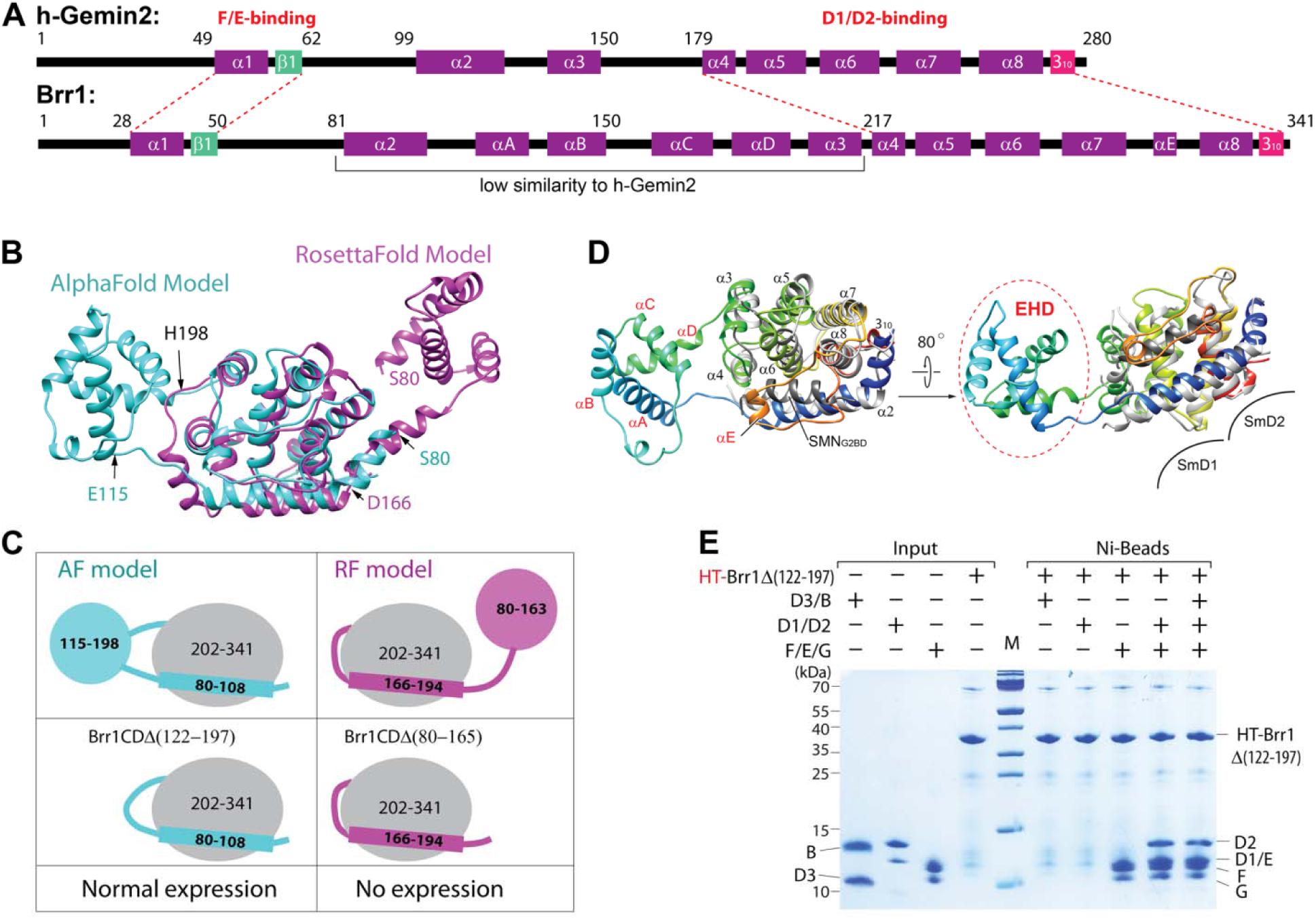
Characterization of the Brr1 structure. (A) Secondary structural elements of human Gemin2 and *S. cerevisiae* Brr1. Known domain compositions are indicated. (B) Superimposition of the AlphaFold and RosettaFold models of Brr1(80-341). (C) The summary of the truncated constructs of Brr1CD and expression test results. See details in Fig. S6. (D) The superimposition of the AlphaFold structure model of Brr1(80-341) (colored in rainbow: the N-terminus in blue, the C-terminus in red) with the structure of human Gemin2-CD (in light gray, PDB: 5XJL). The superimposition reveals that the long loop between α7 and α8 in Brr1 (in orange), which contains αE, occupies the position where SMN_Ge2BD_ (in dark gray) binds to Gemin2. Moreover, the EHD of Brr1 is far away from SmD1/D2. (E) The binding assay was conducted to test the interaction of Brr1Δ(122-197) with each of the three Sm subcomplexes, individually and together. The interaction was assessed using a pull-down assay with Ni-beads. One representative of two independent experiments is shown. M: markers.

When Brr1 is superimposed on Gemin2 in the crystal structure of the SMN(26-62)/Gemin2/D1/D2/F/E/G complex, it is unlikely that the EHD interacts with Sm proteins ^20^ (Fig. 5D). Therefore, we investigated whether the removal of the EHD from full-length Brr1 would still allow it to bind to 5Sm. We generated Brr1Δ(122-197) and found that it expressed well. Importantly, it retained the ability to bind to the Sm subcomplexes similar to the full-length Brr1 (Fig. 5E, compared with Fig. 4A). In addition, the purified EHD could not bind to Brr1Δ(122-197) in a pull-down assay (Fig. S6). Therefore, the EHD is unlikely to play a role in the structure and function of Brr1 and may have been integrated into the coding sequence of the *BRR1* gene during evolution. This is further supported by an *in vivo* assay in which the expression of Brr1Δ(122-197) in the yeast strain *brr1*Δ prevented growth retardation (Fig. 7D, which will be described in detail later).

### Formation of the Lot5/5Sm/Brr1 complex which cannot bind U-snRNA

Building on prior research in vertebrates and *S. pombe* ^5, 12, 18^ along with our biochemical analysis, we hypothesize that Lot5 binds at the opening of the horseshoe-shaped 5Sm, and Brr1 binds at the perimeter. Consequently, the binding of each protein would not sterically hinder the binding of the other. To confirm this hypothesis, we performed GFC analysis. GFC revealed that the mixture of the purified 6S complex with Brr1 eluted earlier (peak at 13.18 ml, aMW=124.9 kDa) than either 6S or Brr1 alone (Fig. 6A), indicating formation of a stable Lot5ΔC/5Sm/Brr1 heteroheptameric complex. Similarly, the mixture of purified Brr1/5Sm with Lot5ΔC also showed earlier elution than either Brr1/5Sm or Lot5ΔC alone, at a position similar to the previous peak (Fig. 6B). It is worth noting that Brr1 showed N-terminal degradation, as confirmed by Western blotting using an anti-FLAG antibody, resulting in the degradation of about 20-25 residues of the N-terminal flexible segment. Nonetheless, this degradation does not affect its ability to bind to 5Sm. Therefore, both Lot5ΔC and Brr1 can bind to 5Sm simultaneously to form a larger 6S/Brr1 complex. These observations align with results from the pull-down analysis of yeast cell extracts (Fig. 1 C-D and Table S1). A pull-down assay corroborated that Lot5ΔC does not bind directly to Brr1 (Fig. 6C), which is in line with the model that the association of Lot5 and Brr1 is mediated by 5Sm. Using less degraded HT-Lot5ΔC to incubate with yeast extract and then conducted Ni-beads pull-down followed by MS analysis, most Sm proteins and Brr1 were unambiguously identified in the Lot5ΔC precipitate (Fig. S7). This evidence further substantiates the presence of the Lot5/5Sm/Brr1 complex within yeast cells.

**Fig. 6.**
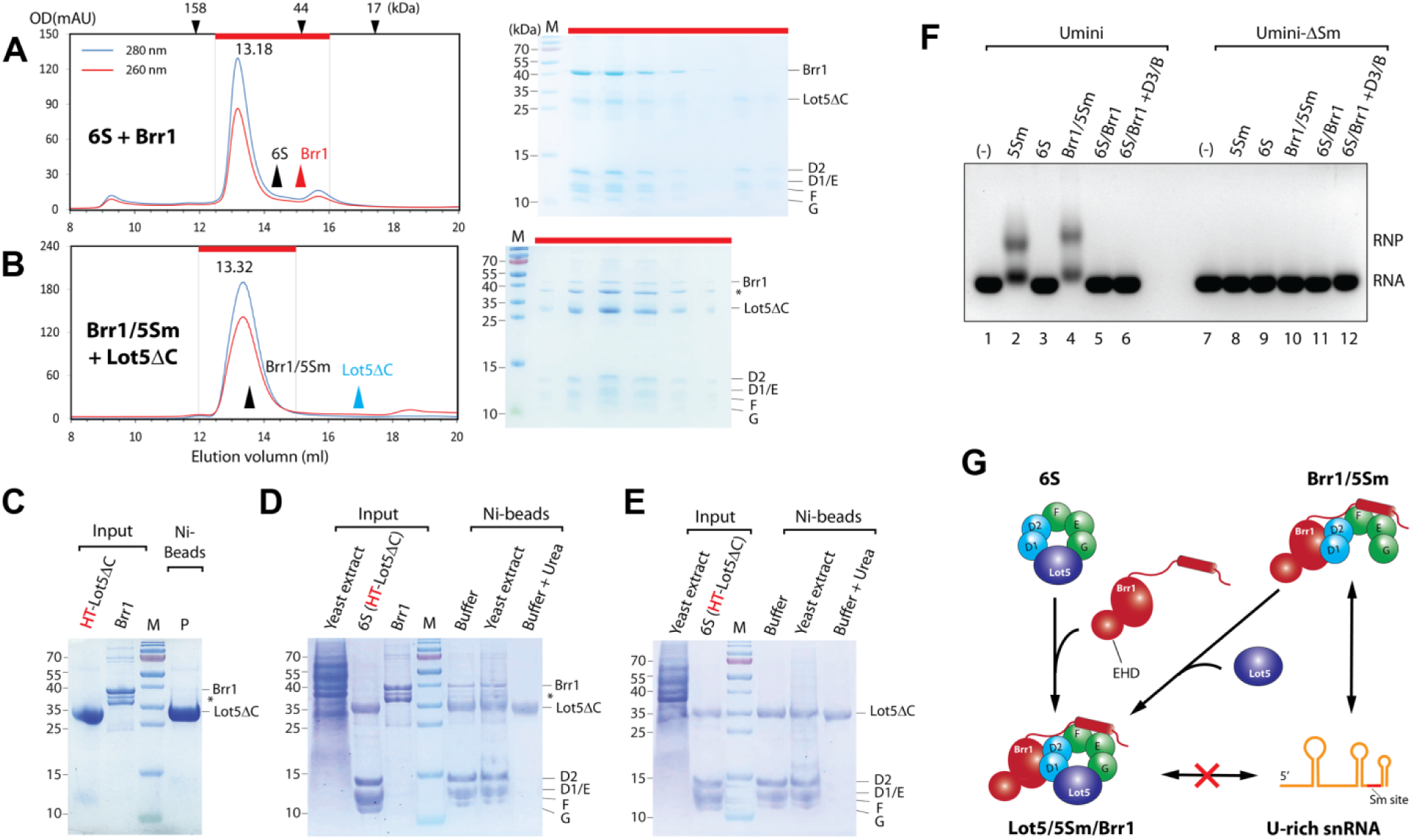
Formation of the Lot5/5Sm/Brr1 complex which inhibits snRNA binding. (A-B) Complexation assays. Preformed 6S mixed with Brr1 (A) or preformed Brr1/5Sm mixed with Lot5ΔC (B) in equimolar amounts, were subjected to GFC (left panels). The eluted fractions (red bar) were then analyzed by SDS-PAGE/CBB staining (right panels). The black and red arrowheads in panel A indicate the GFC peaks of the input components (6S and Brr1) for comparison. The black and blue arrowheads in panel B indicate the GFC peaks of the input components (Brr1/5Sm and Lot5ΔC) for comparison. The asterisk indicates N-terminal degraded Brr1 in panels B-D. (C) No direct interaction between Lot5ΔC and Brr1 tested by Ni-bead pull-down assay. P: precipitate. (D-E) Disassembly assays of 6S/Brr1 (D) or 6S (E) incubated with yeast extract. Pre-assembled 6S/Brr1 or 6S was bound to Ni-beads first, and then incubated with yeast extract (or with washing buffer or washing buffer containing 8M urea as controls). After incubation, the mixtures were separated and washed, and the bound fractions were subjected to SDS-PAGE/CBB staining. Yeast extract input represents 0.5% of total proteins in the reaction. (F) Binding test of different complexes with snRNA using EMSA. Pre-reconstituted 5Sm, 6S, Brr1/5Sm, or 6S/Brr1 or equimolar amounts of 6S/Brr1 mixed with D3/B were pre-incubated with Umini-snRNA or Umini-ΔSm-snRNA (as a control) and subjected to EMSA. M: markers. For each, one representative result of at least two independent experiments is shown. (G) A cartoon model of the formation of the Lot5/5Sm/Brr1 complex and its inhibition of snRNA binding.

To investigate if intracellular factors could dissociate Lot5ΔC from the 6S and 6S/Brr1 complexes, pre-assembled 6S or 6S/Brr1, where only Lot5ΔC bearing an HT-tag, were incubated with yeast extract. The mixtures were subsequently separated with Ni-beads and analyzed by SDS-PAGE/CBB staining. For the control group of either 6S/Brr1 or 6S incubated with washing buffer containing 8M urea, only Lot5ΔC was retained on beads (Fig. 6D-E). In contrast, all 5Sm proteins, Brr1 and Lot5ΔC were retained on beads in the test group for 6S/Brr1 (Fig. 6D), and all 5Sm proteins and Lot5ΔC were retained in the test group for 6S (Fig. 6E), similar to the control groups incubated with only washing buffer. This finding suggests that it is unlikely that there are factors present inside yeast cells that can disassemble the 6S and 6S/Brr1 complexes.

Further analysis using ESMA to examine the binding of 6S/Brr1 or a mixture of 6S/Brr1 and D3/B to Umini-snRNA revealed that, similar to the 6S complex, neither complex could bind Umini-snRNA (Fig. 6F, lanes 5-6). Additionally, neither complex could bind U4-snRNA (Fig. S3). This supports the conclusion that, as shown in Fig. 6G, unlike Brr1/5Sm, the formation of Lot5/5Sm/Brr1, regardless of the order of formation, prevents further assembly of the Sm subcore and core.

### Disruption of genes *LOT5* and *BRR1* in *S. cerevisiae* genome

The analysis outlined above revealed that both Lot5 and Brr1 are involved in binding to Sm proteins and the assembly of the Sm core, which is a critical step in spliceosome biogenesis. We further asked if the deletion of either or both of these proteins in cells could affect the growth of yeast. Using the Cas9-assisted homologous recombination approach, we generated strains *lot5*Δ, *brr1*Δ and *lot5*Δ*/brr1*Δ. These strains were verified by PCR and sequencing and were found to be viable (Fig. 7A and S8). To assess the growth differences, these strains were spotted in serial dilutions on YPD agar and incubated at different temperatures. Strain *lot5*Δ did not display any visible growth retardation compared to the wild-type at temperatures from 16-37°C (Fig. 7B), consistent with previous reports on the disruption of the LOT5 gene ^39^. Strain *brr1*Δ did not show apparent growth difference from the wild-type at 30 and 37°C in terms of colony number at each tested dilution, except for smaller colony sizes at the lower concentrations, but it exhibited slower growth than the wild-type at 16°C (Fig. 7B), consistent with the previous studies ^30, 31^. Interestingly, the *lot5*Δ*/brr1*Δ strain exhibited the same growth pattern as the *brr1*Δ strain at all tested temperatures (Fig. 7B), indicating little or no synergistic effect between the disruption of *BRR1* and *LOT5*. To quantify this observation, we analyzed the growth of these strains in liquid YPD culture at 16 or 37°C (Fig. 7C). At 37°C, all four strains grew at similar rates. However, at 16°C, while *lot5*Δ grew at a rate similar to the wild-type, *brr1*Δ grew slower than both the wild-type and *lot5*Δ strains, and the *lot5*Δ*/brr1*Δ strain grew slightly slower than *brr1*Δ, suggesting more important roles for these factors at low temperatures.

**Fig. 7.**
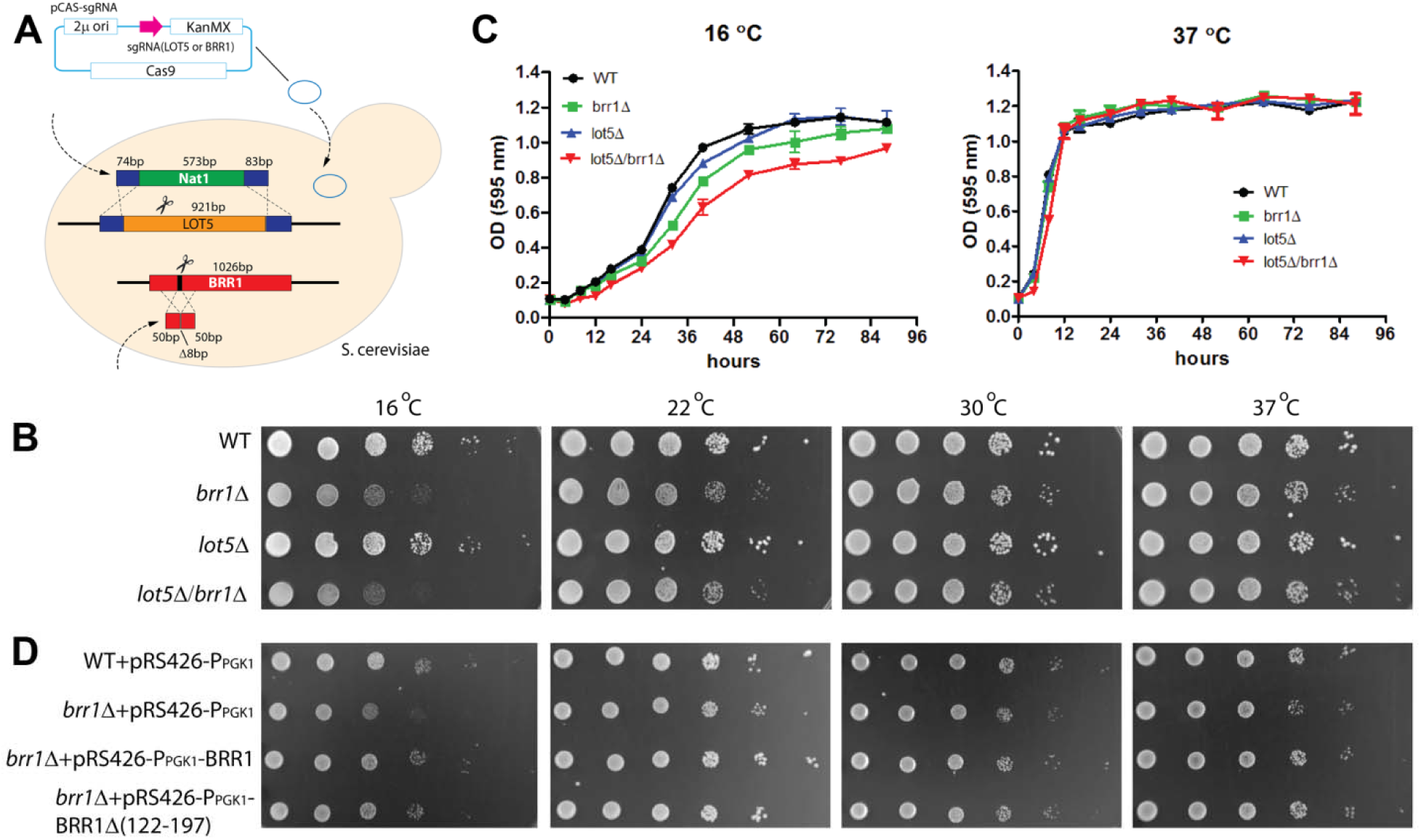
Disruption of genes LOT5 and BRR1 in *S. cerevisiae* causes mild growth retardation. (A) Schematic view of Cas9-assisted disruption of genes LOT5 or/and BRR1 in *S. cerevisiae*. More detailed steps and verifications are in Fig. S8. (B) Growth phenotypes on YPD plates of different strains (*lot5*Δ, *brr1*Δ, and *lot5*Δ/*brr1*Δ) at different temperatures. The tested strains were initially grown in liquid YPD at 30°C to reach an OD_595nm_ of about 0.6. The cell concentrates were then normalized, and serial dilutions of the cells were spotted on YPD plates and cultured at various temperatures (16, 22, 30 and 37°C). The growth of the strains was observed and recorded. (C) Growth time curves in liquid YPD culture at 16 or 37°C. At each time point, samples were collected from three separate cultures for OD_595nm_ measurement. The mean and standard deviation (SD) are shown at each time point. (D) Deletion rescue assay. The *brr1*Δ strain transformed with pRS426-P_PGK1_-BRR1, or pRS426-P_PGK1_-BRR1Δ(122-197) (to test if it functions as full-length Brr1 *in vivo*) or pRS426-P_PGK1_ (as a negative control), or the wildtype strain transformed with pRS426-P_PGK1_ (as a positive control) was spotted in serial dilutions on SD medium plates without uracil and cultured at different temperatures (16, 22, 30 and 37°C). Strain growth was observed and recorded.

In order to investigate the role of *BRR1* in the growth retardation phenotype, we conducted a rescue experiment by expressing BRR1 in the *brr1*Δ strain using the pRS426 plasmid, which contains the URA3 gene. We also transformed the pRS426 vector into the wild-type strain as a positive control, the pRS426 vector into the *brr1*Δ strain as a negative control, and the gene expressing Brr1Δ(122-197), in which the EHD is deleted, into the *brr1*Δ strain for comparison with Brr1. These transformants were spotted in serial dilutions on synthetic dropout (SD) medium without uracil agar and incubated at different temperatures (Fig. 7D). Comparing the *brr1*Δ strain transformed with the pRS426 vector to the wild-type transformed with the same vector, we observed similar cold sensitivity, indicating the vector and the medium components did not contribute to the cold sensitivity of the *brr1*Δ strain. However, when the *brr1*Δ strain was transformed with pRS426-BRR1, we observed a growth phenotype similar to that of the wild-type with the pRS426 vector, indicating a full rescue of the growth retardation at cold temperatures. Interestingly, when the *brr1*Δ strain was transformed with pRS426-BRR1Δ(122-197), the growth retardation at cold temperatures was also rescued. This supports our previous argument that the EHD does not play a significant role in the function of Brr1 inside yeast cells. In summary, our findings suggest that, inside yeast cells, the assembly of the Sm core through a self-interacting (direct) pathway is sufficient for cellular activities, and Brr1 facilitates Sm core assembly, possibly by further stabilizing the intermediate 5Sm complex, which is beneficial for cell survival at low temperatures. However, Lot5 does not seem to play a significant role in the assembly process.

### Overexpression of gene *LOT5* in *S. cerevisiae* inhibits cell growth

Given that Lot5 binds 5Sm or Brr1/5Sm to form 6S or 6S/Brr1 complexes, which are unable to bind snRNA and facilitate Sm core assembly, we hypothesized that overexpression of Lot5 may inhibit the formation of snRNPs and inhibit cell growth, while overexpression of Brr1 may not have the same effect. As expected, overexpression of Brr1 in either the wildtype or *lot5*Δ strain did not affect yeast cell growth (Fig. 8A). However, overexpression of Lot5 in the wildtype strain resulted in slight growth retardation, particularly at 16°C. Remarkably, in the *brr1*Δ strain, overexpression of Lot5 caused significant growth delay at all test temperatures (Fig. 8B). To confirm that the growth inhibition was specifically caused by Lot5 overexpression, the tested strains were selected on YPD medium containing 5-FOA to eliminate the plasmids carrying Lot5 overexpression constructs. Subsequently, the strains were spotted in serial dilutions on YPD plates. We observed that the growth inhibition in both the wildtype and *brr1*Δ strains, which were initially transformed with pRS426-P_PGK1_-LOT5, disappeared (Fig. 8C). This further confirmed that the inhibitory effects on growth were a result of Lot5 overexpression.

**Fig. 8.**
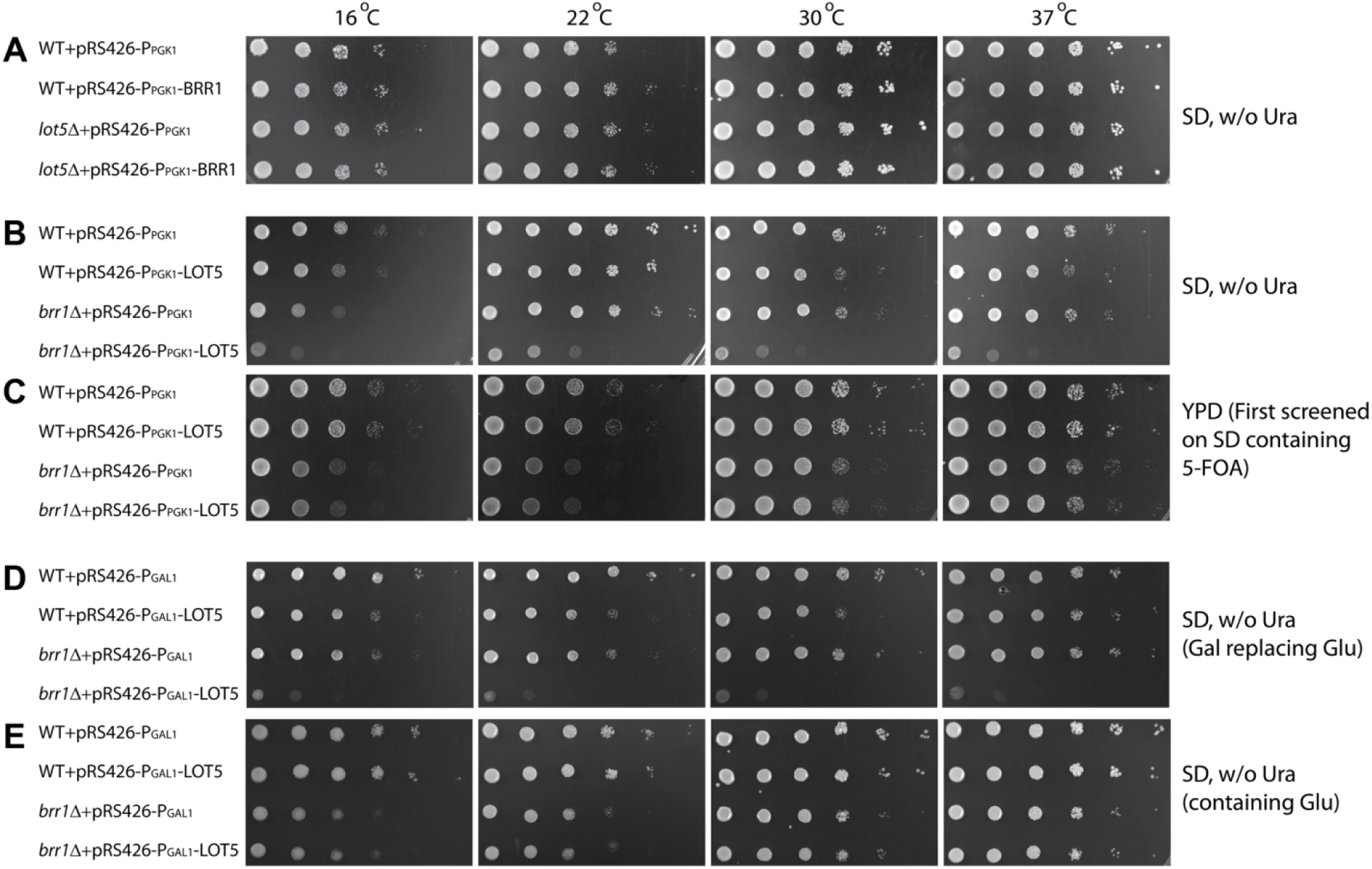
Overexpression of gene LOT5 in *S. cerevisiae* inhibits cell growth. (A) Overexpression of BRR1 has no significant phenotype on cell growth. The wildtype and *lot5*Δ strains transformed with either pRS426-P_PGK1_ (as a negative control) or pRS426-P_PGK1_-BRR1 were spotted in serial dilutions on SD medium plates without uracil and cultured at different temperatures (16, 22, 30 and 37°C). Strain growth was observed and recorded. (B) Constitutive overexpression of LOT5 in the wildtype and *brr1*Δ strains inhibits cell growth. The wildtype and *brr1*Δ strains transformed with either pRS426-P_PGK1_ (as a negative control) or pRS426-P_PGK1_-LOT5 were spotted in serial dilutions on SD medium plates without uracil and cultured at different temperatures (16, 22, 30 and 37°C). Strain growth was observed and recorded. (C) Control experiments for panel B. The four strains from panel B were plated on 5-FOA-containing SD medium to select strains that expelled pRS426 plasmids. Surviving strains were then serially diluted and spotted on YPD plates at different temperatures (16, 22, 30 and 37°C), and their growth was observed and recorded. (D-E) Induced overexpression of LOT5 in the wildtype and *brr1*Δ strains inhibits cell growth. The wildtype and *brr1*Δ strains transformed with either pRS426-P_GAL1_ (as a negative control) or pRS426-P_GAL1_-LOT5 were spotted in serial dilutions on SD medium plates without uracil, with either galactose (D) or glucose (E), and cultured at varying temperatures (16, 22, 30 and 37°C). Strain growth was observed and recorded.

To further confirm the phenotype resulting from Lot5 overexpression in yeast cells, we created an inducible expression plasmid, pRS426-P_GAL1_-LOT5, which contained the inducible GAL1 promoter fused in front of the LOT5 gene. The wildtype or *brr1*Δ strain was transformed with either pRS426-P_GAL1_-LOT5 or pRS426-P_GAL1_ as a control. These transformed strains were grown in SD medium without uracil (containing glucose) until they reached the log phase. Subsequently, they were spotted in serial dilutions on SD medium plates without uracil, where glucose was replaced by galactose to induce Lot5 overexpression. Control plates without uracil were also prepared with glucose as the carbon source.

Similar to the constitutive overexpression of Lot5 mentioned earlier, the induced overexpression of Lot5 resulted in mild growth delay in the wildtype strain. However, in the *brr1*Δ strain, it caused severe growth retardation (Fig. 8D). In contrast, the same strains transformed with pRS426-P_GAL1_-LOT5 or pRS426-P_GAL1_ vector showed no difference in cell growth when glucose was used as the carbon source (Fig. 8E). These observations provide further support that Lot5 and Brr1 have the same roles *in vivo* as observed in biochemical characterization *in vitro*.

## DISCUSSION

In most eukaryotes, from *S. pombe* to vertebrates, the assembly of Sm cores requires multiple assembly chaperones ^1, 4, 5^. However, in *S. cerevisiae*, it was only known that Brr1 plays a role in assembly, yet it is not essential for this process ^30, 31, 40^. This raised the question of whether there are additional assembly chaperones involved in Sm core assembly in *S. cerevisiae* and how they perform their roles. Furthermore, what is the mechanism in *S. cerevisiae* that distinguishes it from other eukaryotes? In this study, we aimed to understand the mechanism of snRNP core assembly in budding yeast and provided answers to these questions. We discovered two pathways in budding yeast, chaperone-mediated and direct assembly pathways, which are distinct from the sole chaperone-mediated pathway in other eukaryotes. The mechanism is summarized in Fig. 9A.

**Fig. 9.**
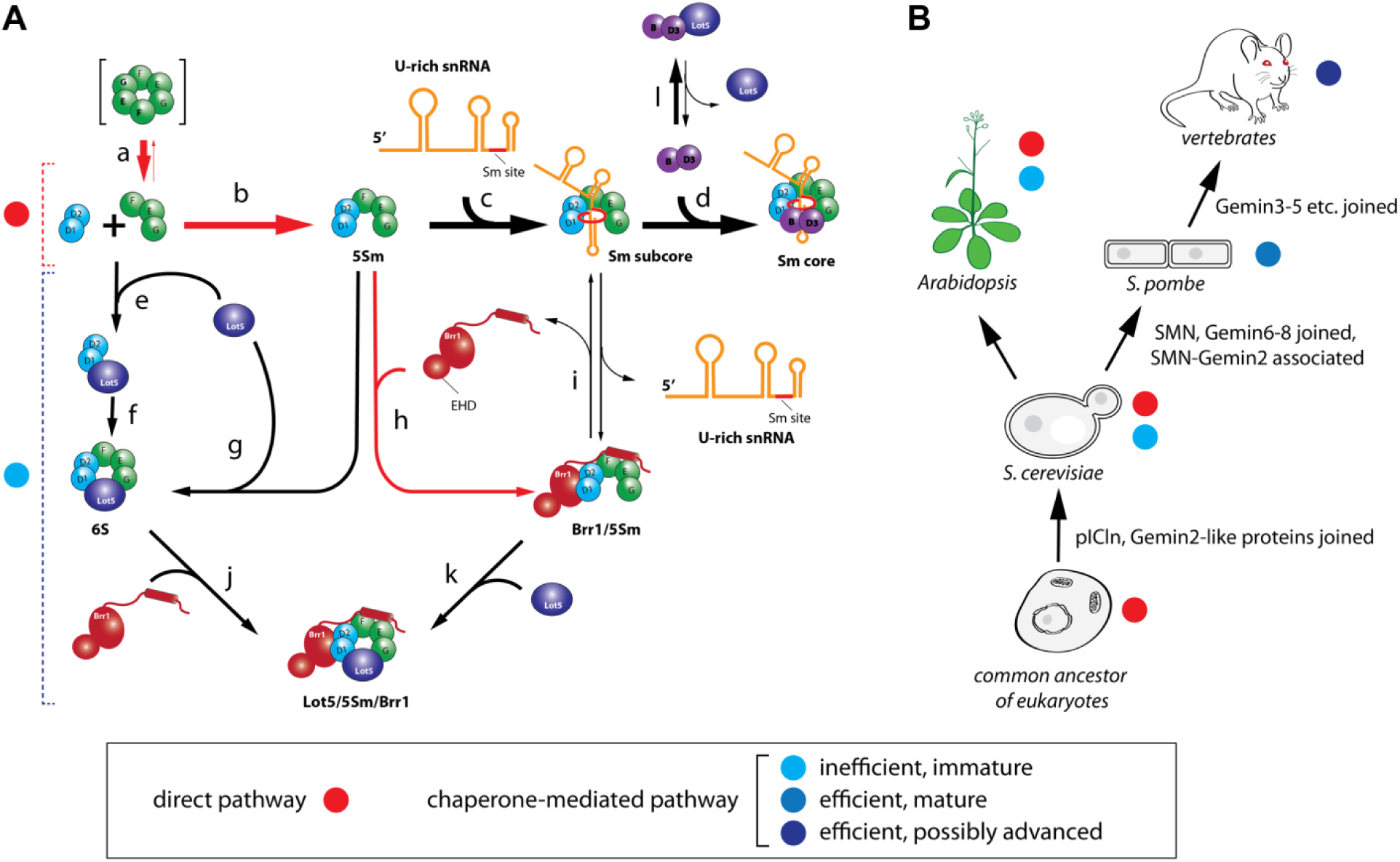
Models of Sm core assembly mechanism in *S. cerevisiae* and evolution of assembly chaperones. (A) The mechanistic model of Sm core assembly in *S. cerevisiae*. There are two pathways for Sm core assembly: (1) the direct assembly pathway, which is dominant; and (2) the chaperone-mediated assembly pathway, which is immature and less efficient compared to the characterized pICln-SMN complex-mediated pathway in *S. pombe* and vertebrates. In the direct pathway, F/E/G exists in a trimeric state (a) so that D1/D2 and F/E/G can spontaneously form a stable 5Sm complex (b). These steps are distinct from those observed in other characterized eukaryotes. This leads to the spontaneous assembly of the Sm subcore (c) and eventually the Sm core (d). In the chaperone-mediated pathway, only the intermediate Brr1/5Sm can progress through the Sm core assembly (h-i), while the intermediate complexes formed with Lot5—either 6S (e, f, g) or Lot5/5Sm/Brr1(j, k)—represent dead-end complexes that are unable to assemble the Sm core. The complexes indicated within square brackets are unstable or transient, and the thickness of the arrow lines represents the preferred direction. The red lines represent reactions unique to *S. cerevisiae* compared to the known mechanisms observed in other eukaryotes. (B) The evolution model of Sm core assembly chaperones. The assembly pathways in *Arabidopsis* and the common ancestor of eukaryotes are anticipated.

In the chaperone-mediated pathway of snRNP core assembly, we discovered that the protein Lot5, whose function was previously unknown, plays a role. It acts as pIClns from vertebrates. It binds to 5Sm, forming a heterohexameric ring (the 6S complex) and simultaneously preventing 5Sm from binding to snRNA. We also found that Brr1, previously considered the homolog of Gemin2, exhibits characteristics that resemble a complex of Gemin2 and SMN_Ge2BD_ rather than acting solely as Gemin2 in its interaction with 5Sm. Brr1 forms a stable complex with 5Sm, whereas individual Gemin2 in most of other species, especially in *S. pombe*, cannot form such a complex. SMN_Ge2BD_ binds to Gemin2 and allosterically enhances its binding to 5Sm ^12^. The Brr1/5Sm complex can accept snRNA, forming the Sm subcore, which further assembles into the Sm core upon joining with D3/B and the simultaneous dissociation of Brr1. By using AI-based Brr1 structural models in combination with biochemical and genetic approaches, we determined that the AF model is more accurate than the RF model. Moreover, we found that the EHD of Brr1 in the AF model is dispensable. Brr1 binds to 6S to form a larger complex Lot5/5Sm/Brr1 in which Lot5 is not displaced. This further indicates that the single protein Brr1, despite being longer than hGemin2 and *Sp*Gemin2 and containing an extra domain, does not function as the holo-SMN complex, which displaces pICln ^5, 12^. Rather, Brr1 merely acts as Gemin2/SMN_Ge2BD_. The binding site of SMN_Ge2BD_ to Gemin2 in other species is likely occupied by a small helix αE in the long loop between α7 and α8 of Brr1, as suggested by both the AF and RF models. Moreover, the preformed Brr1/5Sm complex is also able to accept Lot5 to form the Lot5/5Sm/Brr1 complex. The Lot5/5Sm/Brr1 complex, however, is unable to bind snRNA, representing a dead-end complex in snRNP core assembly. Overall, the chaperone-mediated pathway is less efficient and immature, because the intermediate complexes 6S and Lot5/5Sm/Brr1 cannot bind snRNA and facilitate Sm core formation, although Brr1/5Sm may further stabilize 5Sm and facilitate Sm core assembly. The inhibitory effect of Lot5 on Sm core assembly was further confirmed by overexpression of Lot5 in yeast cells. Overexpression of Lot5 in the wildtype strain caused mild growth retardation, consistent with the result of a previous study involving systematic gene overexpression in *S. cerevisiae* ^41^. Overexpression of Lot5 in the *brr1*Δ strain resulted in significantly slower growth compared to individual Lot5 overexpression or *brr1*Δ alone. This suggests synthetic or synergistic interactions between Lot5 overexpression and loss of Brr1, supporting the notion that Lot5 and Brr1 both function within the same Sm core assembly pathway.

It is noteworthy that in budding yeast, neither Lot5 nor Brr1 is essential for vegetative growth. Deletion of each individual gene has minimal impact on cell growth, and even when both genes are deleted, the cells still exhibit normal growth at 30-37 °C. Only mild growth retardation is observed at cold temperatures (Fig. 7). This is in stark contrast to other eukaryotic species, including unicellular fission yeast and higher vertebrates, where counterparts of Lot5 (such as pICln) and Brr1 (such as Gemin2) play essential roles in Sm core assembly. Additionally, many other assembly factors, such as SMN, Gemin6, Gemin7 and Gemin8, are indispensable for Sm core assembly in these organisms ^7, 8, 9, 10, 12, 13, 14, 15^. The unique dispensability of Lot5 and Brr1 in *S. cerevisiae* suggests that the mechanism for Sm core assembly in budding yeast differs significantly from that of other eukaryotes.

The reason why cells can grow normally particularly at 30-37°C when both Lot5 and Brr1 are disrupted in budding yeast comes from the direct pathway of snRNP core assembly specific to this organism. The stable interaction between D1/D2 and F/E/G and the unique oligomeric state of F/E/G in budding yeast are distinct from those in other eukaryotes. D1/D2 and F/E/G can almost completely form a stable 5Sm heteropentamer, which is significantly different from the counterparts in *S. pombe* (no stable 5Sm forms)^12^ and humans (only ∼50% 5Sm forms) ^35^. Because 5Sm can spontaneously form and be stable enough, there is no need for Lot5 (the fungal equivalent of pICln) to bind D1/D2 and recruit F/E/G to form the heterohexameric 6S complex. In other species, pICln is required to help the formation of 5Sm by forming 6S first, and then the SMN complex displaces pICln from 6S and retains 5Sm on Gemin2 ^5, 12,18^. Surprisingly, in budding yeast, F/E/G exists simply as a heterotrimer, unlike the heterohexameric form seen in *S. pombe* (completely)^12^ and human (in the majority) ^5, 35^. This is likely the main reason why Sc-D1/D2 and Sc-F/E/G can spontaneously form a stable 5Sm, because this reaction does not compete with the dimerization of the F/E/G heterotrimer, as is the case in other species. It is worth noting that although Sm subcomplexes of other eukaryotic organisms such as human and *S. pombe* mixed with snRNAs can spontaneously assemble into Sm cores *in vitro*, the molecular mechanisms are different from that of *S. cerevisiae*. In the former, D1/D2 and F/E/G cannot form a stable 5Sm, which is an energetically uphill reaction. Only when coupled with the energetically downhill reactions of the bindings of snRNAs and D3/B can Sm cores spontaneously form. Inside cells crowded with millions of other proteins, the small amount of 5Sm is probably insufficient to couple with the downstream reactions to assemble enough Sm cores. In contrast, in budding yeast, D1/D2 and F/E/G spontaneously form 5Sm, indicating that this reaction is energetically downhill and each step of Sm core formation is energetically favorable. This enables more efficient spontaneous Sm core assembly even inside cells.

Our study also provides insights into the evolution of snRNP core assembly chaperones in eukaryotes (Fig. 9B). We have found only two proteins, Lot5 and Brr1, involved in Sm core assembly in *S. cerevisiae*, which is a much smaller number than those of other species, such as vertebrates (12 chaperones), and even smaller than that of another unicellular eukaryote, *S. pombe* (6 chaperones). They represent an earlier, more immature version of the assembly chaperone system in evolution compared to the system in *S. pombe*. In *S. cerevisiae*, 5Sm can form spontaneously, and therefore, the Sm subcore and core can assemble sequentially without strict requirements for any assisting proteins. The direct pathway may have been the only pathway for snRNP core assembly in the common ancestor of eukaryotes. This pathway may have facilitated the emergence and evolution of 5Sm-binding proteins, which eventually became the assembly chaperones in high eukaryotes. Interestingly, Lot5 and Brr1 are the proteins that directly bind to the horseshoe-shaped 5Sm at different locations, with one likely binding to the opening and the other to the bulging perimeter, based on the interactions of their homologs with 5Sm ^18, 20^. Lot5 and other pICln homologs belong to the PH domain-like superfamily, one of the largest superfamilies of protein folds ^42, 43^. Although proteins containing this module all consists of a seven-stranded β-sandwich followed by a C-terminal α-helix, they are well known of having very low sequence similarities and playing a wide range of functions ^42^. The fact that PH-like proteins are also identified in bacteria demonstrates that this module existed before the appearance of eukaryotes ^42^. It is plausible that an ancient PH-like protein accumulated a few specific residue mutations in β5 and the C-terminal α-helix, allowing it to gain a new function in interacting with D1/D2, similar to a pICln-like protein. This ancient protein likely gave rise to Lot5 in *S. cerevisiae*. The most complicated domain of Brr1 or Gemin2 orthologues is the C-terminal helical domain which binds D1/D2. The D1/D2-binding regions are located within the conserved α3-α8 fold, which is similar to HEAT-repeat proteins ^44^ and may have derived from an ancient HEAT-repeat protein. The joining of Brr1 in the Sm core assembly possibly occurred before Lot5, because the presence of Lot5 could potentially block Sm core assembly and reduce cell growth, as demonstrated in the overexpression of Lot5 in the *brr1*Δ strain (Fig. 8). In contrast, Brr1 does not inhibit Sm core assembly, but rather stabilizes 5Sm and facilitates Sm core assembly. This suggests that Brr1 evolved to play a role earlier in Sm core assembly, before Lot5. As in the case of budding yeast, the presence of Brr1 in the organism could tolerate the emergence and long-term existence of Lot5, despite Lot5 temporarily having no valuable role in Sm core assembly. Compared to the recently characterized miniature assembly chaperone system (ICln/SMN/Gemin2/Gemin6-8) in *S. pombe* ^12^, Lot5 is similar to ICln, but the single protein Brr1 is less complex and less mature than the 5-membered SMN complex. Brr1 cannot displace Lot5 from 6S, and therefore cannot facilitate Sm core assembly through the Lot5-mediated pathway. Furthermore, there is unlikely an intracellular factor that can displace Lot5 from the 6S and 6S/Brr1 complexes (Fig. 6). Perhaps due to the immature state of Brr1, 5Sm has to form stably and spontaneously in *S. cerevisiae*. During the evolutionary process, with the evolution of SMN, Gemin6-8 and the association between Gemin2 and SMN, a more advanced and mature SMN complex emerged, capable of displacing Lot5 (or pICln) from 5Sm. This upgrade of the chaperone system likely rendered the stable formation of 5Sm and the direct assembly pathway unnecessary. As Sm proteins underwent mutations during evolution, the direct assembly pathway may have been further diminished, leading to increased dependence of the chaperone-mediated pathway for Sm core assembly in most contemporary eukaryotic species. The presence of 5Sm-binding proteins such as Brr1 and Lot5 may have also facilitated mutations in Sm proteins, as demonstrated in previous genetic studies in which the presence of Brr1 tolerated mutations in SmD1, SmD2, SmF and SmG that were synthetically lethal with *brr1*Δ ^30^. In vertebrates, more components, such as Gemin3-5 and Unrip, were added to the SMN complex during evolution, making the chaperone system even more complex. However, the exact role of these members in Sm core assembly in vertebrates remains to be explored.

Interestingly, in *Arabidopsis*, a plant species, there is no SMN orthologue, and Gemin2 is the only component of the SMN-Gemins complex ^29^. Disruptions of the Gemin2 gene in *Arabidopsis* only show mild growth retardation ^29^. Similarly, disruptions of the pICln gene in *Arabidopsis* also lead to only mild growth retardation ^28^. This suggests that the components of the Sm core assembly chaperones and their essentiality in *Arabidopsis* are similar to those in *S. cerevisiae*. It is therefore likely that in *Arabidopsis*, the oligomeric state of SmF/E/G is primarily a trimer, SmD1/D2 and SmF/E/G can spontaneously form a stable 5Sm, and the direct Sm core assembly pathway predominates in these plant cells (Fig. 9B).

SMN deficiency causes SMA in humans and many animal models. In the study of SMA biology, there is still a major question that is not fully understood: why does a universally required SMN protein specifically cause motor neuron degeneration ^17, 45^? Given the discovery of the direct Sm core assembly mechanism in this study, which bypasses the need for any assembly chaperones, including the SMN-Gemins complex, the strategy of direct formation of a stable 5Sm may be employed in genetic engineering in other multicellular species. This strategy could be used to investigate the downstream pathophysiological mechanisms of SMA.

Overall, we have discovered two distinctive pathways in budding yeast for snRNP core assembly: the chaperone-mediated pathway and the direct pathway, distinguishing them from the exclusively chaperone-dependent pathway observed in other eukaryotes. These findings offer valuable insights into the evolutionary dynamics of assembly chaperones across eukaryotes. Additionally, this study has implications for deepening our understanding of SMA-related diseases.

## MATERIALS AND METHODS

### Plasmid constructions for protein expression in *E. coli*

The genes encoding seven Sm proteins, Brr1 and Lot5 were amplified from the *S. cerevisiae* (Sc) genomes extracted using the Yeast genome extraction kit (Tiangen Biotech, China). These Sm genes were then constructed into 3 different plasmids. For SmD1 and SmD2, since the co-expression of full-length SmD1 and SmD2 did not express well, SmD2 and a C–terminal-truncated form of SmD1 (residues 1-109), which keeps the essential Sm fold and removes the nonessential portions for better expression, were used and constructed in the vector pCDFDuet. The plasmid is called pCDFDuet-HT-ScD2-ScD1, which contains a hexameric histidine tag (His_6_) followed by a Tobacco Etch Virus (TEV) cleavage site (His_6_-TEV-tag or HT-tag) at the N-terminal end of SmD2. Similarly, for SmB and SmD3, a truncated form of SmB (residues 1-105) and SmD3 (residues 1-88) were used to improve expression, with the HT-tag at the N-terminal end of SmB. They were constructed in the vector pCDFDuet and named pCDFDuet-HT-ScB-ScD3. Finally, Full-length versions of SmF, SmE and SmG were constructed in the pCDF vector with the HT-tag at the N-terminal ends of both SmE and SmG. The plasmid is named pCDF-ScF-HT-ScE-HT-ScG.

For the Sm core assembly assisting proteins, full-length Lot5, a truncated version of Lot5 (residues 1-227), full-length Brr1, and truncated versions of Brr1 were constructed in the plasmid pET21 with a HT-tag fused at the N-terminus of each protein. In the plasmids, each gene was controlled under a single T7 promoter and contained a ribosomal binding site. To create these constructs, a homologous recombination method called seamless cloning was employed to link the insert fragments and the vector fragments using the Monclone Single Assembly Cloning Mix (Monad Biotech, China). All the created constructs were verified by DNA sequencing.

### Protein expression and purification

The recombinant proteins and protein complexes were all expressed using BL21 (DE3) competent cells. To initiate protein expression, the transformed cells were cultured in Luria-Bertani (LB) medium supplemented with appropriate antibiotics (50 μg/ml), at 37 °C and 200 rpm, until the optical density at 600nm (OD_600nm_) reached about 0.8. To induce protein expression, 0.5 mM isopropyl-D-thiogalactopyranoside (IPTG) was added to the culture, and the cells were allowed to grow overnight at 18 °C and 200 rpm. After the induction period, the cells were collected by centrifugation, and the resulting cell pellet was suspended in lysis buffer [25 mM Tris-HCl, pH 8.0, 250 mM NaCl, 10% glycerol, 20 mM imidazole, 1 mM phenylmethylsulfonyl fluoride (PMSF), 20 mg/L aprotinin, 20 mg/L leupeptin, 20 mg/L pepstatin, 0.2 mM AEBSF and 5 mM β-mercaptoethanol]. The cell suspension was either directly used for purification or stored at - 20°C for use later after snap freezing in liquid nitrogen.

The cells were broken by passing a high-pressure cracker at 700 Bar and 4 °C for 5 minutes, and the supernatant was collected after centrifugation. The overexpressed proteins or protein complexes were purified in general through a series of chromatography steps. First, Ni-NTA chromatography (GE Healthcare, USA) was used for purification. Next, ion exchange chromatography was employed, followed by gel filtration chromatography (GFC) as the final step. In cases where the removal of the His_6_-TEV-tag was needed, TEV protease cleavage was performed after Ni-NTA chromatography. The tag-removed proteins were collected by passing through a Ni-column again at a low concentration of imidazole in the buffer. Throughout the purification process, the quality and quantity of the proteins were monitored by sodium dodecyl sulfate (SDS)-polyacrylamide gel electrophoresis (PAGE) and Coomassie brilliant blue (CBB) staining. Finally, the purified proteins were snap-frozen in liquid nitrogen and stored at - 80°C for future use.

### Protein-protein Interaction by pull-down assay

About 50 μg of Ni-NTA agarose beads were incubated with approximately 0.2 mg of HT-tagged protein or protein complex in the binding buffer (50 mM Tris-HCl, 250 mM NaCl, 20 mM imidazole, 5% glycerol, 0.01% NP-40, and protease inhibitors at pH 8.0) for 1 hour at 4 °C in a small gravity column with both ends of the column sealed. Following the incubation, the beads were washed with 1 ml of binding buffer, and the washing step was repeated five times. Subsequently, about 1.2-fold molar amount of tag-removed protein or protein complex was added to the beads in the binding buffer and incubated at 4 °C for 1 hour with the column ends sealed. After the second incubation, the beads were then washed with 1 ml of binding buffer, and this washing step was again repeated five times. Finally, the proteins bound to the beads were resolved using SDS-PAGE and visualized by CBB staining.

### Protein complexation by GFC

For protein complexation test, the purified proteins or protein complexes were mixed together at an equimolar ratio and incubated at 4 °C for a minimum of 1 hour to allow complex formation. After the incubation, the mixture was centrifuged at 13,000×g at 4 °C for 15 minutes to remove any debris and collect the supernatant. The supernatant was then passed through an analytical gel filtration column called Superdex 200 Increase 10/300 GL (GE Healthcare, USA) with the running buffer (25 mM Tris-HCl, 250 mM NaCl, 1 mM EDTA, pH 7.5). The eluted fractions were collected at intervals of 0.5 ml and analyzed by SDS-PAGE and staining. The apparent molecular weights (aMWs) of the protein complexes were calculated on the basis of the GFC protein standards (Bio-Rad, USA) as a control using the equation: aMW=10^(-0.2331*Vol^ ^+5.1687)^ [R^2^=0.977, aMW(kDa), Vol(ml)]. The theoretical MWs and aMWs of the proteins and complexes in this study can be found in Table S2.

### Western blotting

The protein components separated by SDS-PAGE were transferred to a polyvinylidene fluoride membrane (PVDF) for western blotting. The membrane was washed three times with TBST buffer (25 mM Tris-HCl, 140 mM NaCl, 3 mM KCl, 0.1% Tween-20, pH7.4), and then incubated in a blocking solution (TBST buffer containing 5% nonfat milk) at room temperature for 2 hours. After the incubation, the membrane was incubated with either HRP-conjugated anti-FLAG-M2 monoclonal antibody or HRP-conjugated anti-His-tag monoclonal antibody (both from proteinTech Group, USA) that was diluted to a manufacturer-suggested concentration in the blocking solution at room temperature for 2 hours. After the incubation, the membrane was washed with TBST buffer 3-5 times. The detection of the tagged proteins was performed using an enhanced chemiluminescence (ECL) detection system.

### *In vitro* RNA production and purification

In the Sm core assembly study, we mainly utilized a customized snRNA called Umini-snRNA. Umini-snRNA was created by replacing the noncanonical Sm site with the canonical U4 Sm site in human U7 snRNA. Moreover, three derivatives of Umini-snRNA were used: Umini-ΔSm, in which the UUUUU sequence was replaced with CCCCC, Umini-3’ss, in which the 3’-SL was linearized, and Umini-3’Δ, in which the 3’-SL was completely deleted. Additionally, a wildtype U4-snRNA from *S. cerevisiae* was also included for comparison. Their sequences can be found in Table S3.

All these RNAs were produced using *in vitro* transcription with the T7 High Yield RNA Transcription Kit (Vazyme, China). The DNA templates were created by PCR. The transcribed RNAs were purified using the Spin Column RNA Cleanup & Concentration Kit (Sangon Biotech, China). The RNA quality was assessed using agarose gel electrophoresis, and the concentrations were estimated by measuring their OD_260nm_.

### *In vitro* RNA–protein complex assembly assays

In this study, RNA-protein complex assembly assays were conducted using electrophoresis mobility shift assay (EMSA) and GFC assay. Prior to mixing with proteins, the RNAs were heated to 95°C for 5 minutes and then allowed to cool down at room temperature.

For the EMSA test, a 10 μl reaction mixture was prepared, containing 50 mM Tris-HCl (pH 8.0), 500 mM NaCl, 2.5 mM MgCl_2_, 1 mM ETDA, 0.05 mg/ml heparin, 0.02 nmol of RNA and 0.2 nmol of protein (or protein complex). After incubation at 37°C for 10 minutes, 2 μl of 6× RNA loading buffer was added and the mixture was sampled into a 1% agarose gel. The bands were separated by electrophoresis at 150 V for 15 minutes and visualized using Gel-Red staining. To verify the protein components present in the RNP complex of Umini-snRNA mixed with 5Sm or 5Sm+D3/B, the agarose gel piece containing the RNP band was excised, mixed with sample buffer, and heated at 95°C. Finally, the mixture was analyzed using SDS-PAGE/CBB staining.

For the GFC test, purified protein complexes were incubated with various RNAs in the final volume of 300 μl in an assembly buffer containing 25 mM Tris-HCl, 250 mM NaCl, 2 mM MgCl_2_, and 1 mM EDTA at pH 8.0. The details of the amounts used are described in Table S4. After incubation at 37°C for 30 minutes, the samples were subjected to centrifugation at 15,000 rpm for 20 minutes in a table centrifuge and then applied to a superdex 200 Increase 10/300 GL GFC via a 500 μl sample loop. The eluted fractions were collected in 0.5 ml increments, resolved by SDS-PAGE and visualized using CBB or Gel-Red staining.

### Plasmid constructions for protein expression and genetic disruption in *S. cerevisiae*

For construction of the yeast plasmid p415-TAP-Brr1, p415-TAP-Lot5 or p415-TAP-EGFP, the vector fragment was amplified using PCR from the plasmid p415-GalL-Cas9-CYC1t (Addgene #43804) to keep the GalL promotor and CYC1t terminator but delete the Cas9 sequence which would be replaced by the TAP-tagged protein of interest in the following construction. The TAP fragment, which contained the FLAG-TEV-CBP tag, was subcloned from the plasmid pFLAG-CBP (gift from Dr. Rui Bao, Sichuan University) using PCR. The fragment containing Brr1 or Lot5 was amplified by PCR from yeast genome, and the fragment containing EGFP was amplified from a previous plasmid in our lab. The TAP fragment and the protein fragments were fused together by nested PCR, resulting in a single insert fragment. The insert fragment and the vector fragment were linked using a homologous recombination method called seamless cloning via the Monclone Single Assembly Cloning Mix (Monad Biotech, China).

The gene disruption plasmid pCas-sgRNA (Lot5 or Brr1) was constructed based on the pCAS plasmid (Addgene plasmid # 60847) ^46^. The pCAS plasmid was designed specifically for Cas9-assisted disruption of genes in yeast. It contains a Cas9 gene, a sgRNA cassette, and a KanMX cassette for G418 selection. The sgRNA sequences used for LOT5 and BRR1 were GGCTACCAGATGGTAGGGAA<u>GGG</u> and GTACGATGATGAGGATGAGG<u>GGG</u>, respectively. The underlined three nucleotides are the protospacer adjacent motif (PAM).

The expression plasmids pRS426-P_PGK1_-Lot5 and pRS426-P_PGK1_-Brr1 were utilized for high constitutive expression of Lot5 or Brr1 in yeast. To construct these plasmids, overlapping elongation PCR, was performed to insert the strong constitutive promoter PGK1 upstream of LOT5 or BRR1. The resulting fragments were then cloned into the pRS426 plasmid, which contains a 2μ origin (allowing for high copy number of the plasmid in cells) and a URA3 gene for selection. The plasmids pRS426-P_PGK1_-Brr1 and its derivative pRS426-P_PGK1_-Brr1Δ(122-197) were also used in a deletion rescue assay. Furthermore, for high inducible expression of Lot5 or Brr1 in yeast, the expression plasmids pRS426-P_GAL1_-Lot5 and pRS426-P_GAL1_-Brr1 were constructed using a similar approach, with the inducible promotor GAL1 replacing PGK1. The control vector plasmids pRS426-P_PGK1_ and pRS426-P_GAL1_ were constructed similarly with the promoter sequence directly followed by a stop codon. All the constructed plasmids were verified by DNA sequencing.

### Tandem affinity purification (TAP) from yeast

The yeast plasmid p415-TAP-Brr1, p415-TAP-Lot5 or p415-TAP-EGFP was transformed into yeast strain BY4742 using the LiAc method ^47^. The transformants were plated on synthetic dropout (SD) medium without Leu and incubated at 30°C for 2 days. Colonies were picked and verified by PCR using primers specific to p415 plasmids to ensure the presence of the desired constructs. The verified strains were then inoculated in 1 L of SD liquid medium without Leu and shaken at 30°C for 2 days. A small volume of the samples was used for verification of TAP-fused protein expression by SDS-PAGE and Western blotting (Anti-FLAG M2 monoclonal antibody was used) while the remaining cells were collected after centrifugation and stored for later usage.

The stored cells were re-suspended in a cold lysis buffer (10 mM Tris-HCl, pH 8.0, 40 mM KCl, 150 mM NaCl, 0.2 mM EDTA, 10% Glycerol, 1 mM DTT, 1 mM PMSF and 0.1% Triton-X100) and broken by passing through a high-pressure cracker (1400 Bar, 4 °C) 3-5 times. The supernatant was collected after centrifugation at 15,000 rpm, 4°C, for 1 hour. The TAP-fused proteins were purified using two different affinity columns sequentially. In the first step, an anti-FLAG-affinity gel (Millipore, USA) was used to capture the TAP-fused proteins. After washing with TEV cleavage buffer (10 mM Tris-HCl, pH 8.0,150 mM NaCl, 0.5 mM EDTA,1 mM DTT, 0.1% Triton-X100), the beads were incubated with 10 μg TEV protease in the TEV cleavage buffer at 4°C for 2-8 hours. The pass-through fraction was collected, and the beads were washed using a small volume of the TEV cleavage buffer. The washed factions were pooled with the pass-through fraction. In the second step, the pooled samples from the first step were mixed with Calmodulin Resin (GE healthcare) which had been equilibrated with a CBP buffer (25 mM Tris-HCl, pH 8.0,150 mM NaCl, 1 mM MgAc_2_, 1 mM imidazole, 2 mM CaCl_2_, 0.1% Triton-X100 and 1 mM mercaptoethanol) supplemented with 1.5 mM CaCl_2_. The mixture was incubated at 4°C for 2 hours. After washing with the CBP buffer, the proteins of interest were eluted using a CBP elution buffer (25 mM Tris-HCl, pH 8.0,150 mM NaCl, 1mM MgAc_2_, 1 mM imidazole, 20 mM EGTA, 0.1% Triton-X100 and 1 mM mercaptoethanol). The samples were precipitated by addition of trichloroacetic acid (TCA) to a final concentration of 25% on ice. After centrifugation, the protein pellets were washed twice with cold acetone containing 50 mM HCl and dried in a spin vacuum. The dried pellets were stored for silver staining and mass spectrometry (MS) analysis.

### Disassembly assay of reconstituted complexes with yeast extracts

To assess the disassembly of reconstituted yeast protein complexes, both the 6S and 6S/Brr1 complexes were prepared from purified components, with only Lot5ΔC bearing an HT-tag. For the assembly of 6S, HT-Lot5ΔC (80 μg) and two-fold molar excess of D1/D2 and F/E/G (160 μg each) were mixed with 50 μl Ni-TED beads (BBI, China, Cat# C610030) and incubated at room temperature for 30 minutes. The beads were subsequently washed three times using 1 ml aliquots of a washing buffer (25 mM Tris-HCl, pH 8.0, 500 mM NaCl, 1 mM EDTA, 40 mM imidazole, 5% glycerol, and 0.1% Triton X-100). For the assembly of the 6S/Brr1 complex, an additional 200 μg of Brr1 (a two-fold molar excess) was included during the assembly process which was conducted under identical incubation and wash conditions.

Yeast extract was prepared from a 500 ml YPD culture of the BY4742 strain. Cells were harvested by centrifugation at 4000 rpm for 15 minutes, followed by resuspension in ice-cold washing buffer to yield a final volume of 7 ml. Disruption of cells was carried out through agitating with acid-washed small glass beads in a bead mill device. After centrifugation at 12,000 rpm for 30 minutes, the supernatant was collected and stored on ice for immediate use.

The bead-bound preassembled 6S or 6S/Brr1 complexes were incubated with the prepared yeast extract (7 ml), or equal volumes of washing buffer and washing buffer supplemented with 8M urea as controls. The mixtures were incubated at room temperature for a further 30 minutes. After incubation, the beads were washed three times with 1 ml washing buffer. The bead-retained proteins were then separated by SDS-PAGE and detected via CBB staining.

### Gene disruption of *BRR1* and *LOT5* in *S. cerevisiae*

The haploid strain BY4742 (*MAT*α *his3*Δ*1 leu2*Δ*0 lys2*Δ*0 ura3*Δ*0*) of *S. cerevisiae* was used for the genetic disruption study of the *BRR1* and *LOT5* genes by Cas9-assisted homologous recombination (Fig. S8), which is more efficient than the classical recombination method. For this purpose, a helper plasmid based on pCAS, pCAS-sgRNA (see above), which contains the Cas9-expressing gene and an sgRNA specifically targeting either *BRR1* or *LOT5* gene, was constructed. Donor DNA fragments (Table S5) were generated by PCR. For *lot5*Δ, the donor DNA fragment contained the nourseothricin acetyltransferase 1(*NAT1*) gene flanked by 74 bp and 83 bp homologous sequences to the 5’ and 3’ sides of the *LOT5* gene, respectively. For *brr1*Δ, the donor DNA fragment contained a 100 bp *BRR1* gene sequence, in which an 8 bp fragment including the PAM sequence of the sgRNA-Cas9 targeting sequence was removed. This design introduced a frame-shift disruption in the *BRR1* gene.

For single gene disruption, the corresponding pair of pCAS-sgRNA and donor DNA fragment were transformed into the BY4742 strain using the LiAc method ^47^. For *lot5*Δ, the transformed cells were plated on YPD medium containing 100 μg/ml nourseothricin (NTC) and G418 for selection. For *brr1*Δ, the transformed cells were plated on YPD medium containing 100 μg/ml G418 for growth. A certain number of colonies were picked, and PCR was carried out using specific primer pairs (Table S6) to amplify the genomic fragments, which were then sequenced to select the correct strains. To exclude the pCas-sgRNA plasmids from the yeast cells, both *lot5*Δ and *brr1*Δ strains were grown in YPD liquid medium without antibiotics for multiple generations and then plated on YPD agar medium. Multiple colonies were picked, and each colony was inoculated into YPD liquid medium containing 100 μg/ml G418 for selection. If a colony could not grow in YPD+G418, it was further verified by PCR using a pair of primers specific to the pCas-sgRNA plasmids. The verified *lot5*Δ and *brr1*Δ strains were stored for further use.

To generate the *lot5*Δ/*brr1*Δ double gene disruption strain, while avoiding potential lethal phenotypes, the Lot5-expressing plasmid pRS426-P_PGK1_-Lot5, which also contains a URA3 gene, was first transformed into the *lot5*Δ strain to establish the *lot5*Δ+pRS426-P_PGK1_-Lot5 strain, which was selected on SD medium without uracil (Ura) plates. Subsequently, the pair of pCAS-sgRNA and the donor DNA fragment for *brr1*Δ were transformed into the *lot5*Δ+pRS426-P_PGK1_-Lot5 strain. The transformants were plated on SD medium without Ura containing 100 μg/ml of G418 for growth. After selecting colonies with disruption of the *BRR1* gene as described in the single gene disruption protocol above, the *lot5*Δ /*brr1*Δ+pRS426-P_PGK1_-Lot5 strain was plated on SD medium without Ura supplemented with 0.1% 5-fluorolactic acid (5-FOA) and 50 μg/ml Ura. The cells that were able to grow under this selection pressure were considered to have established the *lot5*Δ /*brr1*Δ strain. The pCAS-sgRNA plasmid was also excluded in the same way as described above. All the single and double gene disruption strains used in growth characterization experiments were confirmed to be free of plasmids inside the cells.

Stains BY4742 (wildtype), *lot5*Δ, *brr1*Δ and *lot5*Δ /*brr1*Δ were inoculated in liquid YPD and grown at 30°C until they reached an OD_595nm_ of about 0.6. After normalizing the cell concentrates, the cells were spotted in serial 1:10 dilutions on YPD plates. These plates were then cultured at four different temperatures (16, 22, 30 and 37°C) for 2-5 days. Moreover, for making growth curves, all the strains were inoculated into liquid YPD with the same initial concentration and cultured at both 16 and 37°C for 3-4 days. Each strain was triplicated, and small volumes were taken at different time points to measure OD_595nm_ values.

### Deletion rescue assay in *S. cerevisiae*

For the *brr1*Δ strain, deletion rescue assay was performed by transformation of the plasmid pRS426-P_PGK1_-Brr1 and its derivative pRS426-P_PGK1_-Brr1Δ(122-197). Transformation of pRS426-P_PGK1_ was carried out as a negative control, while the wildtype strain transformed with pRS426-P_PGK1_ served as a positive control. The transformants were selected by plating on SD medium plates without Ura. Once established, the strains were selected and cultured in liquid SD medium without Ura at 30°C until they reached an OD_595nm_ of approximately 0.6. Following concentration normalization, the cells were spotted onto SD medium plates without Ura in serial 1:10 dilutions. The plates were then incubated at four different temperatures (16, 22, 30 and 37°C) for 2-5 days, and the growth of the strain was observed and recorded.

### Overexpression of Lot5 or Brr1 in *S. cerevisiae*

For the overexpression of Brr1, the constitutive expression plasmid pRS426-P_PGK1_-Brr1 or the control plasmid pRS426-P_PGK1_ was transformed into the wildtype and *lot5*Δ stains. The transformants were plated on SD medium without Ura for selection. Once established, the strains were picked and grown in liquid SD medium without Ura at 30°C until they reached an OD_595nm_ of about 0.6. After normalizing the cell concentrations, the cells were spotted in serial 1:10 dilutions onto SD medium plates without Ura.

Regarding Lot5 overexpression, two approaches were taken: constitutive and inducible overexpression. For constitutive Lot5 overexpression, similar to Brr1, the constitutive expression plasmid pRS426-P_PGK1_-Lot5 or the control plasmid pRS426-P_PGK1_ was transformed into the wildtype and *brr1*Δ stains. The strain selection and growth assay were identical to those for Brr1. Due to significant phenotypic changes observed in Lot5 overexpression, additional verification to determine the relationship between the phenotype and plasmid was conducted. To do so, the plasmids were excluded from the test strains by plating them on SD medium supplemented with 0.1% 5-FOA and 50 μg/ml Ura. The resulting plasmid-free strains were then spotted in serial 1:10 dilutions onto YPD plates for comparison.

For inducible Lot5 overexpression, the inducible expression plasmid pRS426-P_GAL1_-Lot5 or the control plasmid pRS426-P_GAL1_ was transformed into the wildtype and *brr1*Δ stains. The transformants were plated on SD medium without Ura for selection. Once established, the strains were picked and grown in liquid SD medium without Ura at 30°C until they reached an OD_595nm_ of about 0.6. After normalizing the cell concentrations, the cells were spotted in serial 1:10 dilutions onto two types of plates: (1) SD medium plates without Ura, in which galactose replaced glucose for induced expression, and (2) classic SD medium plates without Ura, which contained glucose and served as a control.

All these plates were cultured at four different temperatures (16, 22, 30 and 37°C) for 2-5 days, and the growth of the strain was observed and recorded.

### Protein identification by mass spectrometry (MS)

The preparation of protein samples, including in-gel digestion by trypsin and peptide extraction, followed our previous method in general ^48^. The peptide samples were sent to the MS facility of Western China Hospital for liquid chromatography (LC)/MS/MS analysis. The LC and MS/MS devices used were the ThermoFisher Vanquish Neo and ThermoFisher Orbitrap Exploris 480, respectively. Peptide mass information was searched against the S. cerevisiae proteomic database to generate a list of potentially matched proteins. The list was further refined based on a combination of the molecular weights of the pulldown proteins on SDS-PAGE, unique peptide number, coverage of unique peptides, and score.

### Structure models of the proteins

The structure models of Lot5 and Brr1 (Fig. S1) were obtained from two different sources. For Lot5, the model was directly downloaded from the AlphaFold protein structure database (https://alphafold.ebi.ac.uk/) ^49^. For Brr1, two structure models were obtained: (1) from the AlphaFold protein structure database, and (2) from the Rosetta Online Server (https://rosie.rosettacommons.org/) after submission of the sequence of Brr1. Additionally, the RosettaFold model of the Brr1-CD (residues 80-341) region (Fig. S1) was also obtained from the same server and was almost identical to the corresponding region in the full-length structure model.

### Sequence alignments

Sequence alignment of Lot5 with pICln homologs from various species was conducted using the MAFFT server (https://mafft.cbrc.jp/alignment/server/). The structural alignment of the AlphaFold structure model of Lot5 against the fruit fly pICln crystal structure (PDB code: 4F7U) was utilized as a constraint. The alignment was further refined by Clustal Omega in the JalView 2.11 program ^50^. Similarly, the sequence alignment of Brr1 with Gemin2 homologs from different species was performed using the MAFFT server. The structural alignment of the AlphaFold structure model of Brr1 against the human Gemin2 crystal structure (PDB code: 5XJL) was used as a constraint. The alignment was then optimized using Clustal Omega in the JalView 2.11 program ^50^.

## AUTHOR CONTRIBUTIONS

Y. Wang, X. Chen, X. Kong and Y. Chen performed the interaction and complexation studies of the proteins and protein complexes. Y. Wang and X. Chen performed the interaction studies of the proteins with RNAs. Y. Wang and X. Kong performed the over-expressions and genetic disruptions in *S. cerevisiae*. Y. Wang, X. Kong, Y. Chen and Z. Xiang performed structural studies of Brr1. Y. Xiang, Y. Hu, Y. Hou, S. Zhou, C. Shen, L. Mu and D. Su participated in this project. Y. Wang, X. Chen and X. Kong helped in preparation of the paper. R. Zhang conceived, designed and supervised the project, and wrote the paper.

## DATA AND MATERIALS AVAILABILITY

All data needed to evaluate the conclusions in the paper are present in the paper and/or the Supplementary Materials.

