## supplemental material 1 for "A unique mechanism of snRNP core assembly"

Yingzhi Wang *et al.*

**This PDF file includes:**

Tables S1 to S6

Figs. S1 to S8

**Other Supplementary Materials for this manuscript include the following:**

Data S1

**Table S1. Mass spectrometry identification of the interacting proteins of Brr1 or Lot5.** Although the initial proteomic analysis against the *Saccharomyces cerevisiae* protein database revealed hundreds of potential matching proteins, we refined this list by considering the Mascot score, molecular weight (MW), and the band intensity observed in SDS-PAGE coupled with silver staining (Fig. 1C&D). Despite the refinement, the list still included numerous proteins typically abundant in cells, such as ribosomal proteins, ubiquitin, histone proteins, and cytochrome c. Yet, among these, the proteins Brr1, Lot5, and several Sm proteins stood out as the most likely candidates for interacting with our tagged proteins and were selected for subsequent detailed characterization.

| | | **Brr1 pull-down** | | | | **Lot5 pull-down** | | | |
|---|---|---|---|---|---|---|---|---|---|
| **Name** | **#AAs** | **Coverage (%)** | **#PSM** | **# UP** | **Score Mascot** | **Coverage (%)** | **#PSM** | **# UP** | **Score Mascot** |
| **Brr1** | 341 | 52.49 | 123 | 26 | 2870.2 | 43.99 | 35 | 20 | **840.6** |
| **Lot5** | 306 | 16.34 | 12 | 4 | **322.2** | 33.99 | 160 | 9 | 2998.9 |
| **SmD1** | 146 | 66.44 | 55 | 13 | 686.0 | 41.10 | 8 | 6 | 156.1 |
| **SmD2** | 110 | 68.18 | 47 | 8 | 865.2 | 48.18 | 6 | 5 | 86.2 |
| **SmF** | 86 | 41.86 | 18 | 4 | 361.0 | 9.30 | 1 | 1 | 33.5 |
| **SmE** | 94 | 22.34 | 4 | 2 | 32.4 | ND | ND | ND | ND |
| **SmG** | 77 | 22.08 | 2 | 2 | 58.7 | 11.69 | 1 | 1 | 44.5 |
| **SmD3** | 101 | 49.50 | 4 | 4 | 86.4 | 41.58 | 5 | 3 | 110.0 |
| **SmB** | 196 | 50 | 15 | 10 | 380.7 | 22.45 | 6 | 4 | 145.8 |

#AAs: the number of amino acids; #PSM: peptide spectrum match number; #UP: unique peptide number.
ND: not detected.

**Table S2. The proteins and complexes used in this study.**

| Component | Length (residues) | MW (kDa) | GFC Elution volumn (ml) | Apparent MW (kDa) |
|---|---|---|---|---|
| SmD1 | 109 | 12.0 | NA | NA |
| HT-SmD2 (SmD2)[#] | 127 (111) | 14.9 (12.9) | NA | NA |
| HT-SmE (SmE)[#] | 115 (95) | 12.8 (10.4) | NA | NA |
| SmF | 86 | 9.7 | NA | NA |
| HT-SmG (SmG)[#] | 108 (78) | 12.4 (8.5) | NA | NA |
| HT-SmB (SmB)[#] | 126 (107) | 14.5 (12.0) | NA | NA |
| SmD3 | 88 | 9.7 | NA | NA |
| HT-FL-Lot5 | 337 | 38.2 | NA | NA |
| HT-Lot5ΔC | 258 | 29.7 | 16.9 | 17.0 |
| HT-Brr1 (Brr1)[#] | 379 (342) | 44.3 (39.9) | 15.1 | 44.6 |
| HT-Brr1CDΔ(122-197) | 236 | 27.5 | NA | NA |
| HT-Brr1CDΔ(80-165) | 225 | 26.1 | NA | NA |
| HT-Brr1(116-197) | 131 | 14.9 | NA | NA |
| HT-Brr1Δ(122-197) | 314 | 36.3 | NA | NA |
| Brr1Δ(122-197) | 266 | 31.1 | NA | NA |
| D1/D2 | 220 | 24.9 | 16.8 | 17.9 |
| F/E/G | 259 | 28.6 | 16.0 | 27.5 |
| D3/B | 195 | 21.8 | 16.8 | 17.9 |
| 5Sm* | 479 | 53.5 | 15.1 | 44.6 |
| HT-Lot5ΔC/D1/D2 | 478 | 54.6 | 14.9 | 49.6 |
| HT-Lot5ΔC/D3/B | 453 | 51.5 | 16.0 | 27.5 |
| 6S* | 737 | 83.2 | 14.5 | 61.5 |
| HT-Brr1/F/E/G | 649 | 73.8 | 13.6 | 99.7 |
| HT-Brr1/5Sm | 869 | 98.7 | 13.54 | 102.9 |
| HT-Brr1/5Sm/HT-Lot5ΔC | 1127 | 128.4 | 13.18 | 124.9 |

[#]the tag-removed proteins and their lengths and MWs are in brackets.

*5Sm: D1/D2/F/E/G; 6S: HT-Lot5ΔC/D1/D2/F/E/G.

NA: not applied.

**Table S3. The RNA sequences used in this study.**

| Name | RNA sequence |
|---|---|
| Umini | GGGAAGUGUUACAGCUCUUUUAGAAUUUUUGGAGUAGGCUUU CUGGCUUUUCACCGGAAAGCCCCU |
| Umini-ΔSm | GGGAAGUGUUACAGCUCUUUUAGAACCCCCGGAGUAGGCUUU CUGGCUUUUCACCGGAAAGCCCCU |
| Umini-3’ss | GGGAAGUGUUACAGCUCUUUUAGAAUUUUUGGAGUAGGCUUU CUGGCUUUUCAGGCCUUUCGGGGA |
| Umini-3’Δ | GGGAAGUGUUACAGCUCUUUUAGAAUUUUUGG |
| U4 (S. cerevisiae) | GGAUCCUUAUGCACGGGAAAUACGCAUAUCAGUGAGGAUUCG UCCGAGAUUGUGUUUUUGCUGGUUGAAAUUUAAUUAUAAACC AGACCGUCUCCUCAUGGUCAAUUCGGUGUUCGCUUUUGAAUA CUUCAAGACUAUGUAAAUUUUUGGAAUACCUUU |

The Sm site is highlighted in red and the ΔSm sequence in blue. The sequences underlined are the strands forming the double-stranded helical “stem” of 3’-stem-loop.

**Table S4. The amounts and ratios of RNAs and proteins used in GFC studies.**

| components | amount（molar ratio） |
|---|---|
| Lot5ΔC | 500 μg |
| Brr1 | 500 μg |
| F/E/G | 1 mg |
| D1/D2 | 500 μg |
| D3/B | 500 μg |
| F/E/G+D1/D2 | 250 μg+250 μg (1:1) |
| Brr1+F/E/G | 600 μg+500 μg (1:1) |
| Brr1+5Sm | 400 μg+500 μg (1:1) |
| Lot5ΔC+D3/B | 500 μg+500 μg (1:1) |
| Lot5ΔC +D1/D2 | 500 μg+500 μg (1:1) |
| Lot5ΔC +5Sm | 250 μg+450 μg (1:1) |
| Brr1/5Sm+Lot5ΔC | 750 μg+225 μg (1:1) |
| 6S+Brr1 | 500 μg+250 μg (1:1) |
| 5Sm+D3/B | 500 μg+250 μg (1:1) |
| Umini-snRNA | 1 mg |
| Umini-snRNA +5Sm | 200 μg+400 μg (1:1) |
| Umini-snRNA +5Sm+D3/B | 200 μg+400 μg+200 μg (1:1:1) |
| 6S + Umini-snRNA | 600 μg+200 μg (1:1) |
| 6S + Umini-snRNA +D3/B | 300 μg+100 μg+200 μg (1:1:1) |
| 6S +D3/B | 600 μg+400 μg (1:1) |

In the table the names and used amounts of RNAs are labeled in blue.

6S: Lot5ΔC/5Sm

**Table S5. The DNA sequences used to disrupt the LOT5 and BRR1 genes in *S. cerevisiae* genome.**

| Name | DNA sequence |
|---|---|
| DNA-Nat1-for-LOT5 | AAGATAGACCATAGCTACAGTGATAGACGAATATTTATTTCAAAAGAAG<br>CATCAACCATAAAATAAAAAGAAAAatgggtaccactcttgacgacacggcttaccggtaccg<br>caccagtgtcccgggggacgccgaggccatcgaggcactggatgggtccttcaccaccgacaccgtcttcc<br>gcgtcaccgccaccggggacggcttcaccctgcgggaggtgccggtggacccgcccctgaccaaggtgttc<br>cccgacgacgaatcggacgacgaatcggacgacggggaggacggcgacccggactcccggacgttcgtc<br>gcgtacggggacgacggcgacctggcgggcttcgtggtcgtctcgtactccggctggaaccgccggctgacc<br>gtcgaggacatcgaggtcgccccggagcaccgggggcacggggtcgggcgcgcgttgatggggctcgcg<br>acggagttcgcccgcgagcggggcgccgggcacctctggctggaggtcaccaacgtcaacgcaccggcg<br>atccacgcgtaccggcggatggggttcaccctctgcggcctggacaccgccctgtacgacggcaccgcctcg<br>gacggcgagcaggcgctctacatgagcatgccctgcccctaaACAACATCTATTATGGGAACA<br>TCCCGCTGTACTATGCGGTCTCGTCCTCTACGAATATGGCTATTTGCCTTC<br>GTATATACCTT |
| DNA-Δ8bp-for-BRR1 | TCTGCAAAAGAGGACGCGTCACAAATCAAGCATGTACGATGATGAGGATG<br>CTTAAAAGGCACGCCATCTCGCCATCCTTGATTAGGCTTCAAAGGAATGT |

The Nat1 gene is in small case and highlighted in green. The 5'-side homolog sequences of both DNA segments are highlighted in yellow. The 3'-side homolog sequences are highlighted in cyan.

The DNA-Nat1-for LOT5 was made by a nested PCR method. Briefly, we first used PCR to generate three DNA segments containing the 5'-side homolog part, Nat1 gene and 3'-side homolog part with the primers overlapping the neighboring sequences (See Table S6). Then we used the mixture of all the three PCR products as a template and the primer pair at the edges (Lot5-HR-F1 and Lot5-HR-R2) to run PCR again to produce the final donor DNA fragment.

**Table S6. The primers used in construction of plasmids and DNA donors, and verification of disruption of genes in *S. cerevisiae*.**

| Purpose | Primer name | Primer sequence |
|---|---|---|
| Making pRS426-$P_{PGK1}$-LOT5 | PGK-Lot5-F | AACAAAAGCTGGAGCTCGTCAGGCATGAACGCATCACAGAC |
| | PGK-Lot5-R | CATTGTTTTATATTTGTTGTAAAAAGTAGATAATTACTTCCTTG |
| | Lot5-F | TACAACAAATATAAAACAATGATGAAAAAAAAACCAAAGTGCCAAATTG |
| | Lot5-R | AGCCGGATCTCACTCGAGTCATTCATTGTCTCTGGAGTTCTTTCTGCCAG |
| Making pRS426-$P_{PGK1}$-BRR1 | PGK-Brr1-F | TCACTATAGGGCGAATTGTCAGGCATGAACGCATCACAGAC |
| | PGK-Brr1-R | TCTTTTCATTGTTTTATATTTGTTGTAAAAAGTAGATAATTACTTCCTTGATGATC |
| | Brr1-F | TCACTATAGGGCGAATTGTCAGGCATGAACGCATCACAGAC |
| | Brr1-R | ACTAATTACATGATTATTCTATCAAGTCTTTTTGACCATAGTTCACTGC |
| Making pCAS-sgRNA (for *brr1Δ*) | CasB-F | TCCTGAGAAGGCAGAAGAGAGTTTTAGAGCTAGAAATAGCAAGTTAAAATAAGGCTAG |
| | CasB-R | CTCTTCTGCCTTCTCAGGAAAAGTCCCATTCGCCACCCGAAG |
| Making pCAS-sgRNA (for *lot5Δ*) | CasL-F | GGCTACCAGATGGTAGGGAAGTTTTAGAGCTAGAAATAGCAAGTTAAAATAAGGCTAG |
| | CasL-R | TTCCCTACCATCTGGTAGCCAAAGTCCCATTCGCCACCCGAAG |
| Making DNA-Δ8bp-for-BRR1 | Brr1-HR-F | TCTGCAAAAGAGGACGCGTCACAAATCAAGCATGTACGATGATGAGGATGCTTAAAAGG |
| | Brr1-HR-R | ACATTCCTTTGAAGCCTAATCAAGGATGGCGAGATGGCGTGCCTTTTAAGCATCCTCAT |
| Making DNA-Nat1-for-LOT5 | Lot5-HR-F1 | AAGATAGACCATAGCTACAGTGATAGACGAATATTTATTTCAAAAGAAGCATCAACCAT |
| | Lot5-HR-R1 | CGTGTCGTCAAGAGTGGTACCCATTTTTCTTTTTATTTTATGGTTGATGCTTCTTTTG |
| | Lot5-HR-F2 | GAGCATGCCCTGCCCCTAAACAACATCTATTATGGGAACATCCCGCTGTACTATG |
| | Lot5-HR-R2 | AAGGTATATACGAAGGCAAATAGCCATATTCGTAGAGGACGAGACCGCATAGTACAGCG |
| Verification of *lot5Δ* | P1-F | CACCATACTTCAACACAGGCCAAGATAGAC |
| | P1-R | CGAAGGCAAATAGCCATATTCGTAGAG |
| Verification of *brr1Δ* | P2-F | GTTCGCTTTGGATAATGTATCATACAATAACATTGC |
| | P2-R | GTAAGAATGAGTCAATCAAAGTAATAACACTTTTTAC |

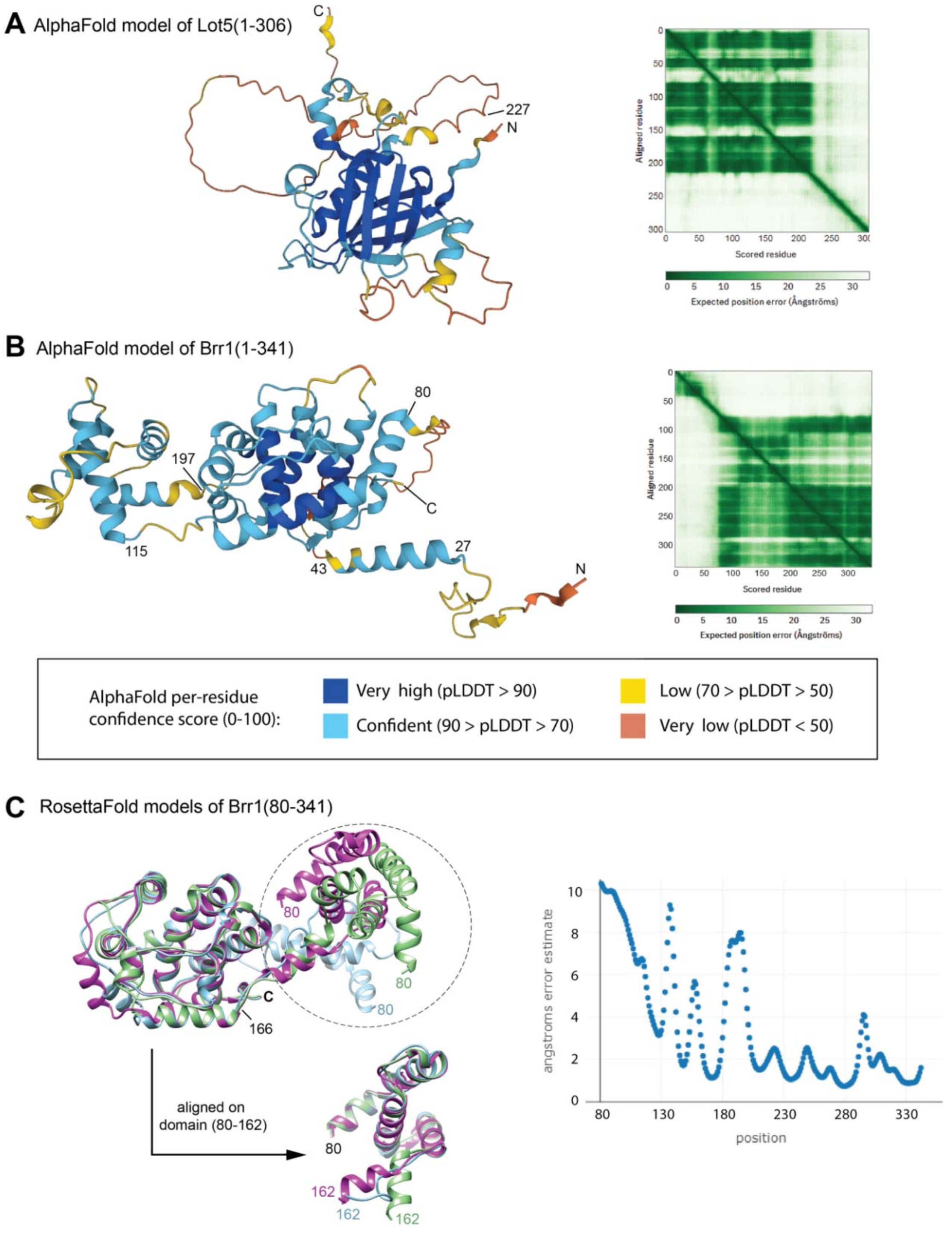


**Fig. S1. The models of Lot5 and Brr1 generated by AlphaFold (https://alphafold.ebi.ac.uk/) and RosettaFold (https://rosie.rosettacommons.org/).** (A-B) The AlphaFold structural models of Lot5 (A) and Brr1 (B) are shown in cartoon representation and colored according to confidence score (left); Expected position error was plotted against residues (right). (C) The top 3 RossettaFold models of Brr1 (residues 80-341), superimposed on the domain (166-341) (left). Expected position error for model 1 (in purple, which is also used in Fig. 5B) was plotted against residues (right).

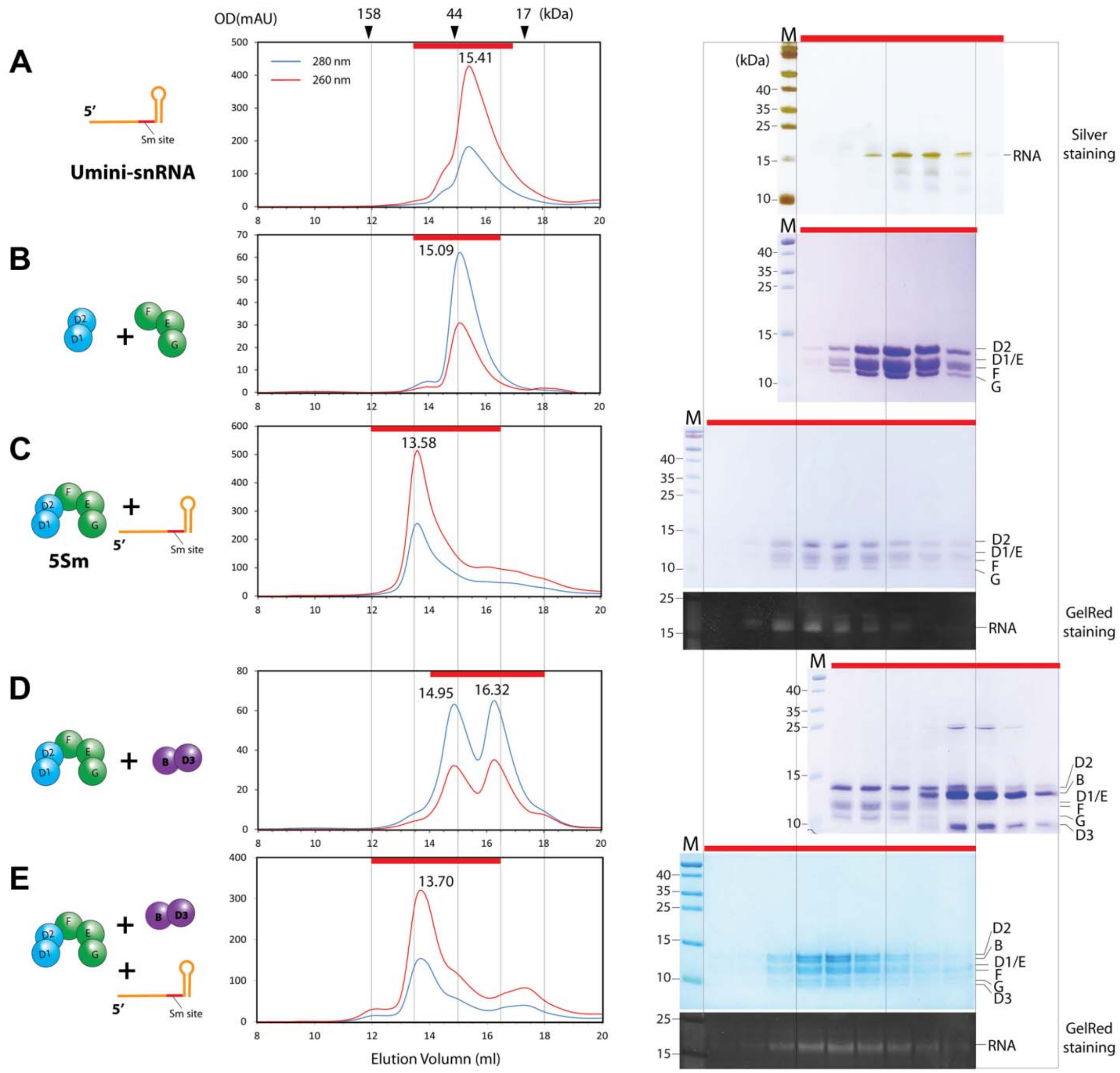


**Fig. S2. Sm subcore and core assembly using Umini-snRNA, tested by GFC.** (A) *In vitro* synthesized Umini-snRNA (left, in cartoon) was subjected to GFC analysis (middle) followed by SDS-PAGE and silver-staining (right). (B-E) Mixture of equimolar amounts of D1/D2 and F/E/G (B), 5Sm and Umini-snRNA (C), 5Sm and D3/B (D) or 5Sm and D3/B and Umini-snRNA (E) (left, in cartoon) was subjected to GFC (middle). The elution fractions, collected each 0.5 ml, were subjected to SDS-PAGE followed by CBB staining or GelRed staining (right). For each, one representative result from at least two independent experiments was shown. Panel B is the same as Fig. 2F and shown here for easy comparison with panel C.

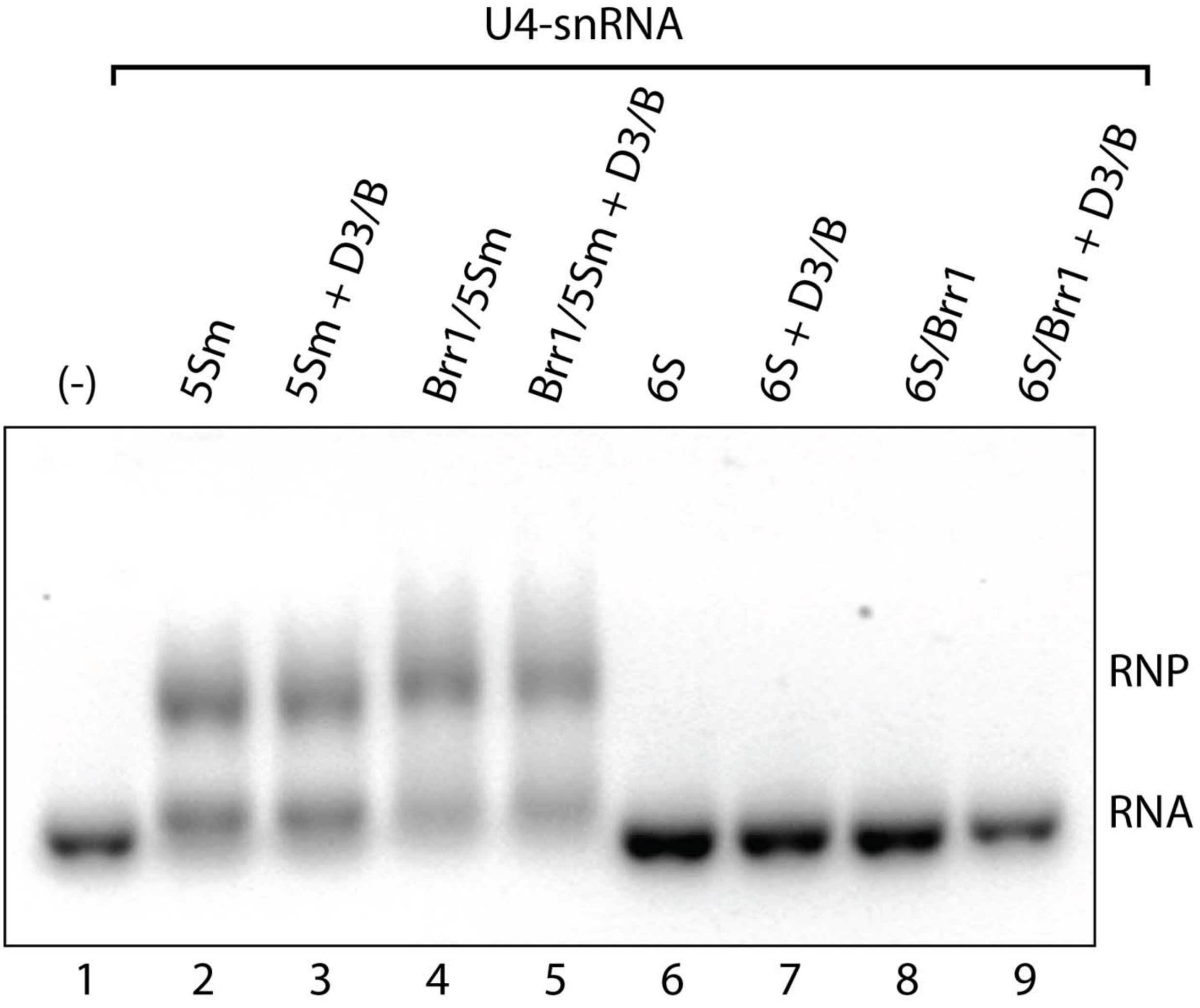


**Fig. S3. The effects of Brr1 and Lot5Δ in Sm subcore and core formation using wildtype *S. cerevisiae* U4-snRNA.** Preformed 5Sm, Brr1/5Sm, 6S, or 6S/Brr1, either alone or mixed with D3/B, was pre-incubated with U4-snRNA from *S. cerevisiae* and subjected to electrophoresis mobility shift assay (EMSA). One representative from two independent experiments is shown.

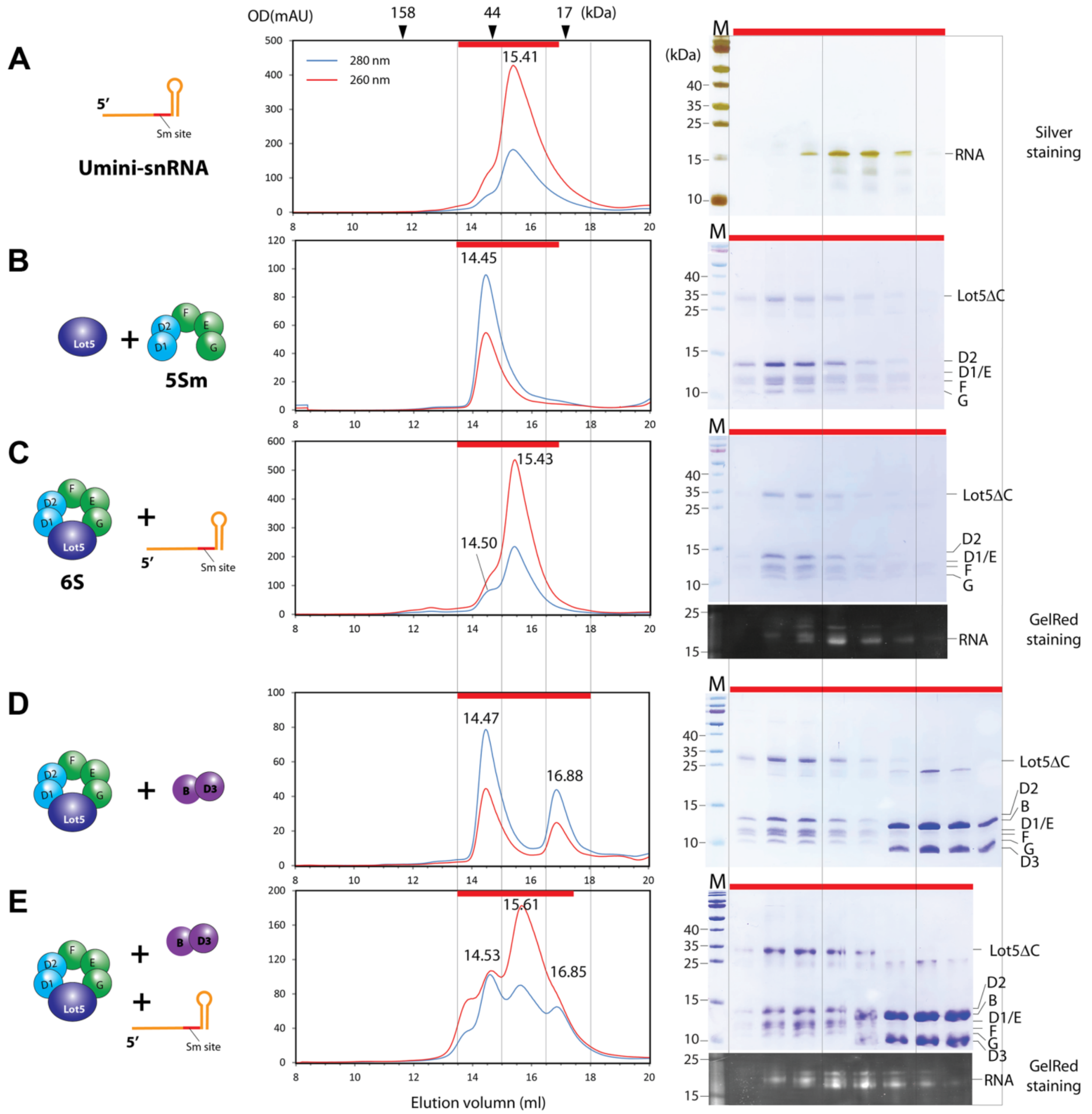


**Fig. S4. Formation of 6S by interaction between Lot5ΔC and 5Sm blocks the assembly of snRNA into the Sm subcore and core, tested by GFC.** (A) *In vitro* synthesized Umini-snRNA (left, in cartoon) was subjected to GFC analysis (middle) followed by SDS-PAGE and silver-staining (right). Panel A is the same as Fig. S2A and shown here for easy comparison with panels C and E. (B-E) Lot5ΔC blocks Sm subcore and core formation. Mixture of equimolar amounts of Lot5ΔC and 5Sm (B), 6S and Umini-snRNA (C), 6S and D3/B (D) or 6S and D3/B and Umini-snRNA (E) (left, in cartoon) was subjected to GFC (middle). The elution fractions, collected each 0.5 ml, were subjected to SDS-PAGE followed by CBB staining or GelRed staining (right). For each, one representative result from at least two independent experiments was shown. Panel B is the same as Fig. 3F and shown here for easy comparison with panel C.

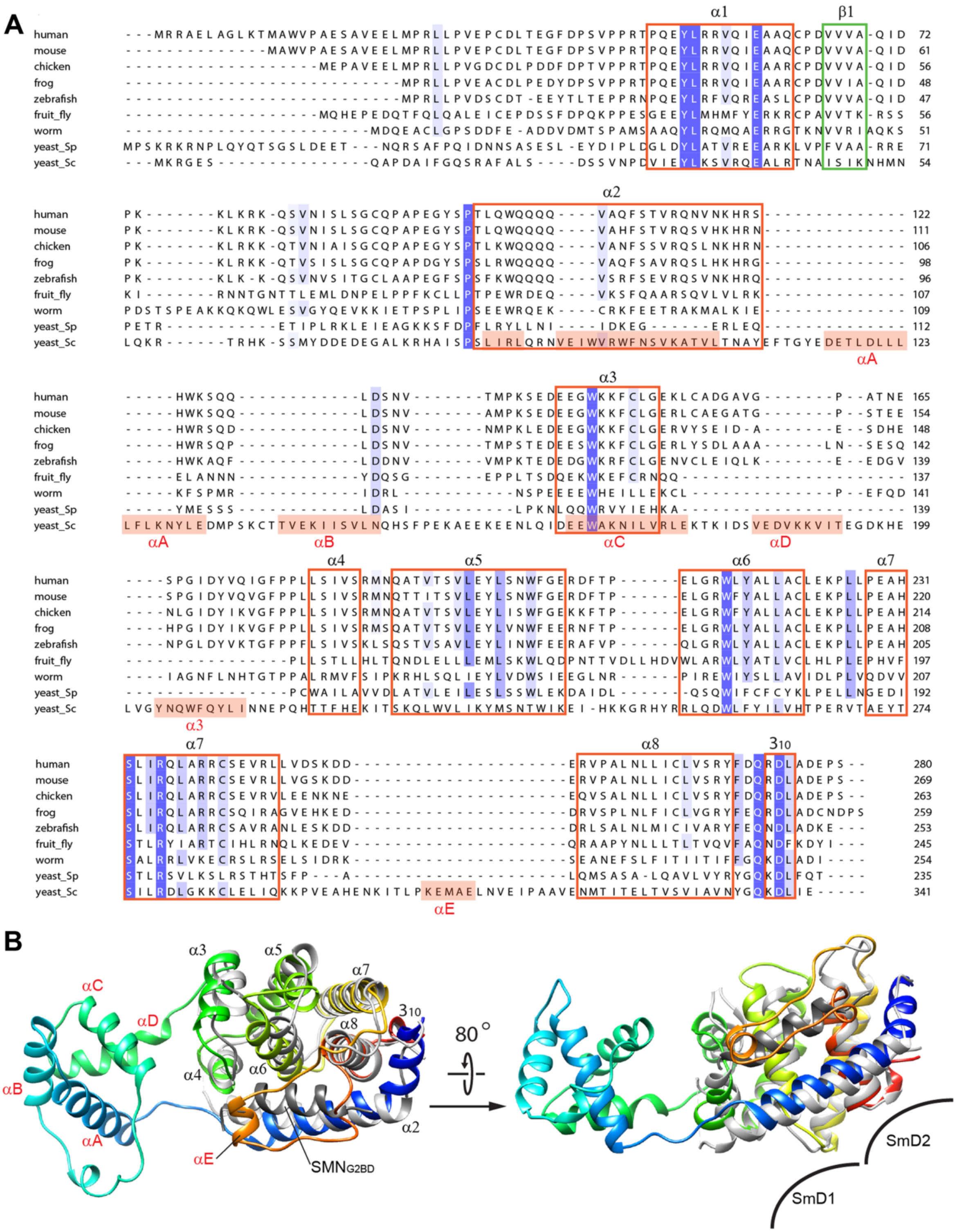


**Fig. S5. Sequence and structure alignments of Brr1 with other homologs.** (A) Sequence alignment of Brr1 (yeast_Sc) with the Gemin2 homologs from various species. The SmF/E-binding domain contains α1 and β1, and the D1/D2-binding domain contains α5-α8 and the last $3_{10}$ helix. (B) Superimposition of the AlphaFold structure model of Brr1(80-341) (colored in rainbow) with human Gemin2-CD (in gray, PDB: 5XJL). Panel B is the same as Fig. 5D and shown here for easy comparison with panel A.

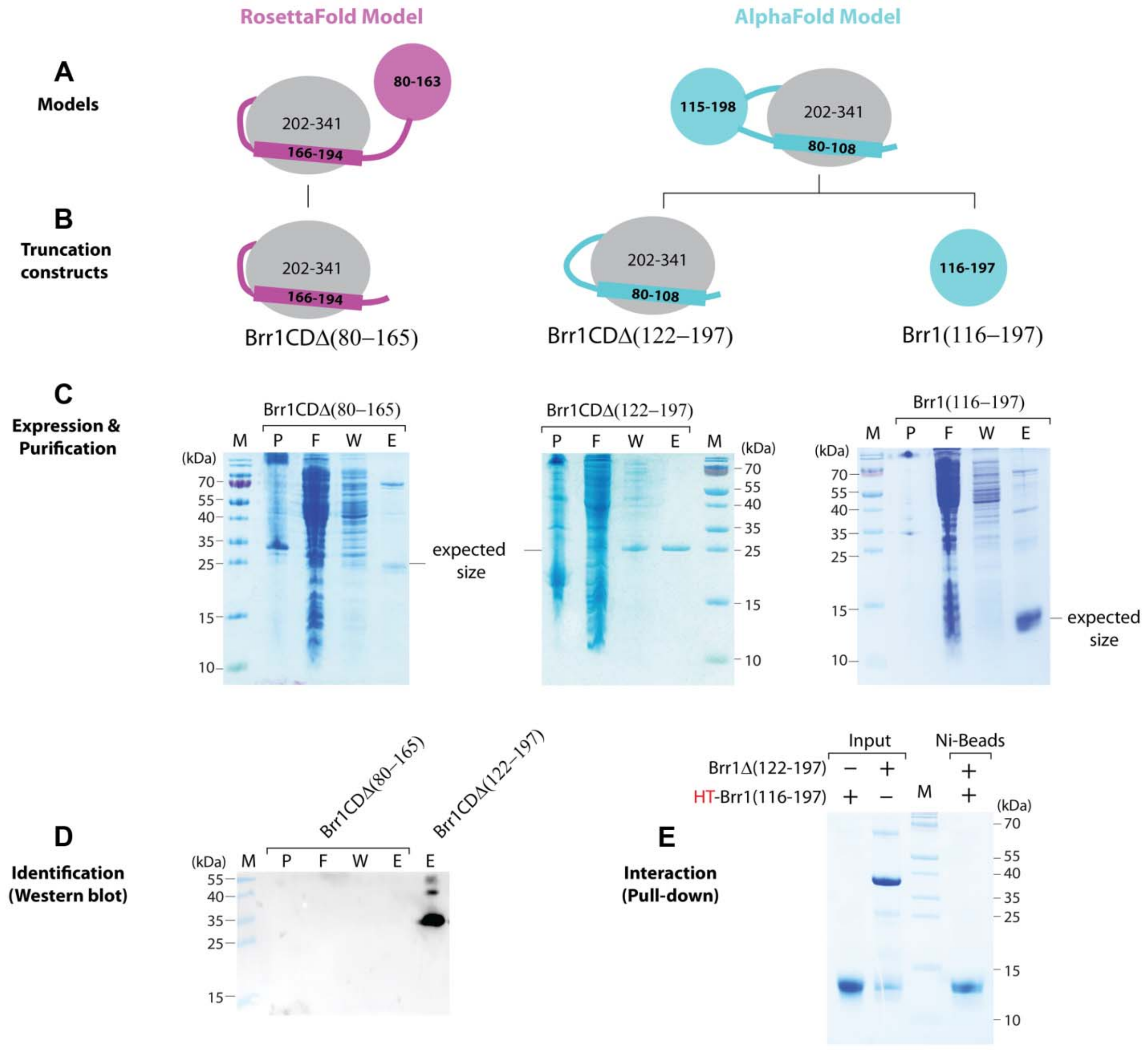


**Fig. S6. Verification of the structure models of Brr1-CD.** (A) Two models are shown in cartoon. (B) Construction of truncated Brr1 proteins (in cartoon). (C) Purification of the truncated Brr1 proteins expressed in *E. coli* by Ni-beads. (D) The samples were further probed by anti-His antibody. (E) Interaction test between Brr1(116-197) and Brr1Δ(122-197) by Ni-beads pull-down assay. M: markers; P: pellet; F: flow-through; W: wash; E: elution.

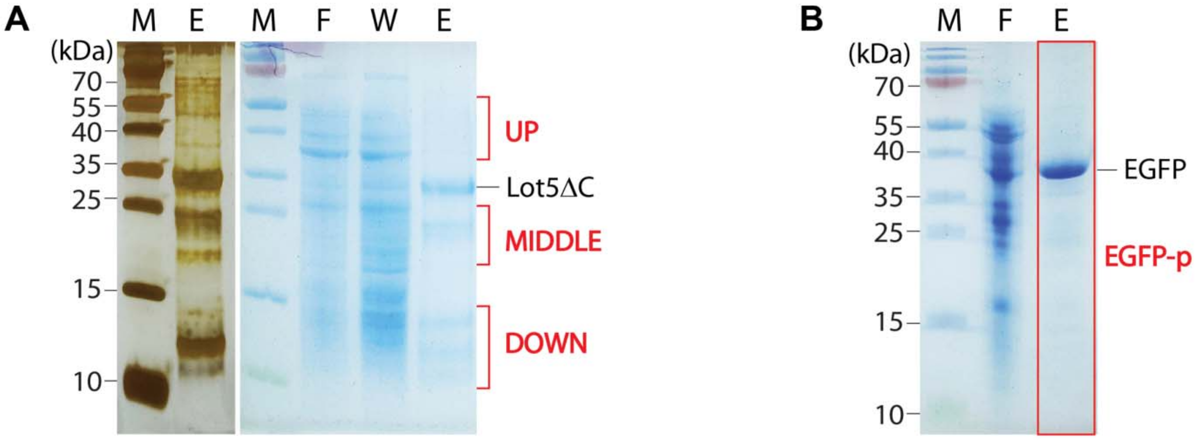


C

| Section | Protein information | | | Lot5ΔC-pulldown | | | EGFP-pulldown | | |
|---|---|---|---|---|---|---|---|---|---|
| | Protein ID | Protein name | MW(kDa) | Intensity | Unique peptides | Unique sequence coverage(%) | Intensity | Unique peptides | Unique sequence coverage(%) |
| UP (38-55kDa) | Q99177 | Brr1 | 39.859 | 1.13E9 | 8 | 28.7 | 4.97E6 | 1 | 2.6 |
| MIDDLE (20-26kDa) | P40018 | SmB | 22.379 | 2.74E7 | 3 | 16.8 | 6.15E6 | 2 | 14.3 |
| DOWN (9-17kDa) | Q06217 | SmD2 | 12.853 | 9.92E8 | 6 | 48.2 | 1.48E7 | 1 | 10.9 |
| | Q12330 | SmE | 10.375 | 6.94E8 | 3 | 43.6 | — | — | — |
| | P54999 | SmF | 9.6599 | 1.99E8 | 3 | 39.5 | — | — | — |
| | P43321 | SmD3 | 11.224 | 1.28E7 | 1 | 8.9 | — | — | — |
| EGFP-p | | EGFP | 32.248 | — | — | — | 1.07E9 | 12 | 46.4 |

**Fig. S7. Identification of the interacting proteins with Lot5ΔC in yeast extract.** (A) Purified HT-Lot5ΔC (about 500 µg) was mixed with yeast extract (from 250 ml YPD culture) in 4 ml of binding buffer (25 mM Tris-HCl, pH 8.0, 250 mM NaCl, 1 mM EDTA, 40 mM imidazole, 5% glycerol, and 0.1% Triton X-100) and the mixture was incubated at 16°C overnight. After incubation, 0.5 ml Ni-beads was added into the mixture for additional 30 minute incubation. The Ni-beads were then washed multiple time with the binding buffer and eluted with 3 ml of elution buffer (25 mM Tris-HCl, pH 8.0, 250 mM NaCl, 1 mM EDTA, 300 mM imidazole, 5% glycerol, and 0.1% Triton X-100). The samples in each fraction were analyzed by SDS-PAGE followed by CBB and silver staining. Three sections of the elution lane, UP, MIDDLE, and DOWN, were cut out and sent to MS facility for protein identification. (B) Purified HT-EGFP (about 500 µg) served as a negative control and was conducted using a similar procedure to HT-Lot5ΔC in panel A. The entire elution lane on SDS-PAGE was cut out for MS analysis. (C) Identification of main interacting proteins with Lot5ΔC. MW range of the cut gel, intensity of MS signal, unique peptide number, and unique peptide coverage were used to select the potential interacting proteins. Brr1 and Sm proteins were unambiguously identified as the interacting proteins with Lot5ΔC. M: markers; F: flow-through; W: wash; E: elution. EGFP-p: EGFP precipitate.

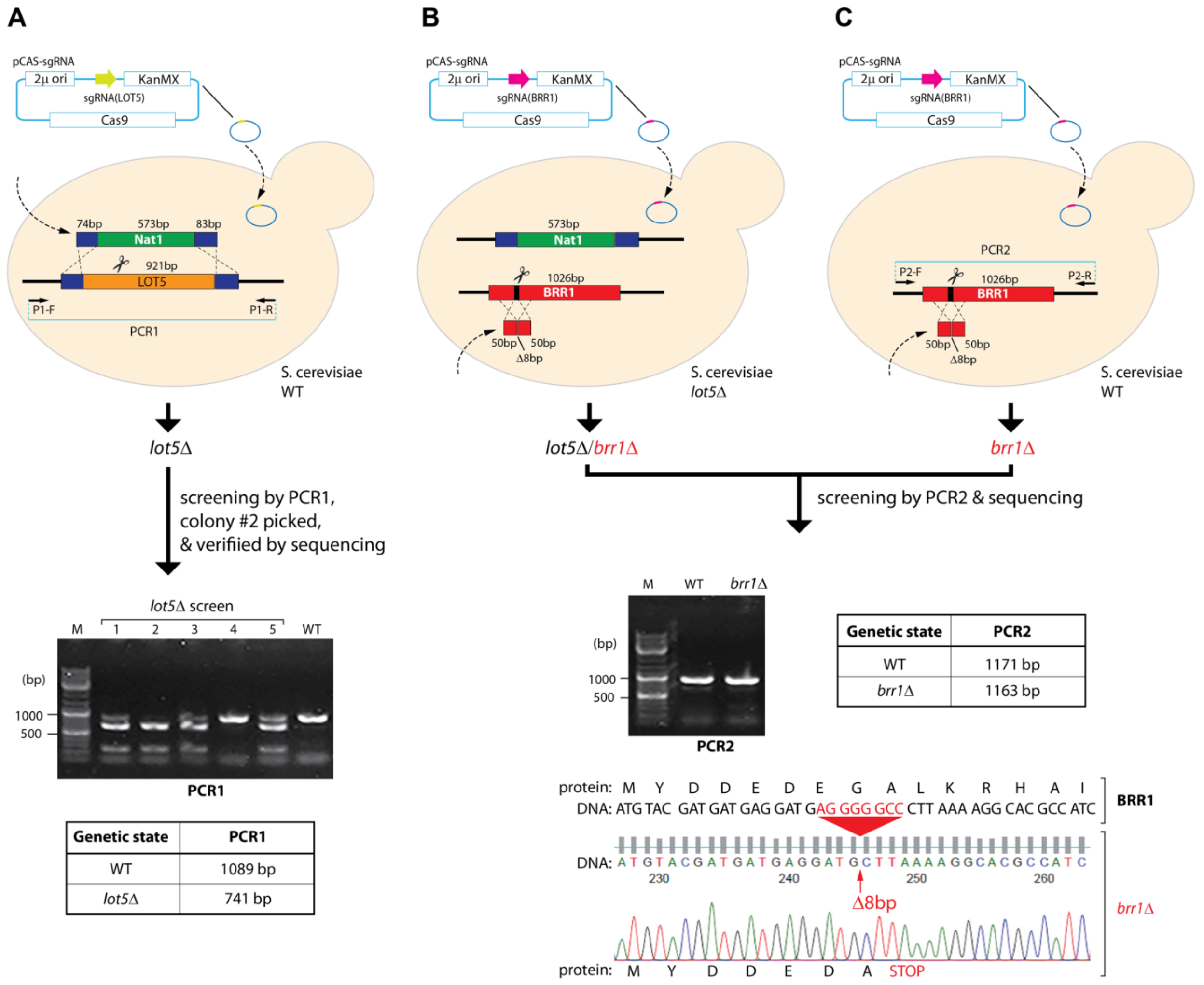


**Fig. S8. The strategies of gene disruptions and verifications of the genotypes.** (A) Construction of strain *lot5Δ* by Cas9-assisted homologous recombination. The strain was screened by PCR and sequencing. (B-C) Constructions of strain *lot5Δ/brr1Δ* (B) and strain *brr1Δ* (C) by Cas9-assisted homologous recombination, based on strain *lot5Δ* and wild-type respectively. Due to the small difference between the sizes of the designed frameshift disruption and wild-type BRR1 gene, the correct genotypes were screened by amplification of the segment of interest by PCR and DNA sequencing. One successful sequenced example is shown. WT: wild-type.
